# Large-scale visualisation of α-synuclein oligomers in Parkinson’s disease brain tissue

**DOI:** 10.1101/2024.02.17.580698

**Authors:** Rebecca Andrews, Bin Fu, Christina E. Toomey, Jonathan C. Breiter, Joanne Lachica, Ru Tian, Joseph S. Beckwith, Lisa-Maria Needham, Gregory J. Chant, Camille Loiseau, Angèle Deconfin, Kenza Baspin, Peter J. Magill, Zane Jaunmuktane, Oliver J. Freeman, Benjamin J. M. Taylor, John Hardy, Tammaryn Lashley, Mina Ryten, Michele Vendruscolo, Nicholas W. Wood, Lucien E. Weiss, Sonia Gandhi, Steven F. Lee

## Abstract

Parkinson’s disease (PD) is a common neurodegenerative condition characterised by the presence in the brain of large intraneuronal aggregates, known as Lewy bodies and Lewy neurites, containing fibrillar α-synuclein. According to the amyloid hypothesis, these large end-stage species form from smaller soluble protein assemblies, often termed oligomers, which are proposed as early drivers of pathogenesis. To date, however, this hypothesis has remained controversial, at least in part because it has not been possible to directly visualise oligomeric aggregates in human brain tissue. Therefore, their presence, abundance and distributions have remained elusive. Here, we present ASA-PD (Advanced Sensing of Aggregates - Parkinson’s Disease), an imaging method to generate large-scale α-synuclein oligomer maps in post-mortem human brain tissue. We combined autofluorescence suppression with single-molecule fluorescence methods, which together, enable the detection of nanoscale α-synuclein aggregates. To demonstrate the utility of this platform, we captured ∼1.2 million oligomers from the anterior cingulate cortex in human post-mortem brain samples from PD and healthy control patients. Our data revealed a specific subpopulation of nanoscale oligomers that represent an early hallmark of the proteinopathy that underlies PD. We anticipate that quantitative information about oligomer distributions provided by ASA-PD will enable mechanistic studies to reveal the pathological processes caused by α-synuclein aggregation.

## Introduction

Parkinson’s disease (PD) is a progressive neurodegenerative disorder that initially causes the loss of dopaminergic neurons in the substantia nigra, resulting in a movement disorder consisting of tremors, bradykinesia, and rigidity.^1^ The disease spreads over several years, affecting multiple brain regions and resulting in dementia, neuropsychiatric, autonomic and sleep disturbances.^2^ Pathologically, PD is characterised by neuronal loss accompanied by the accumulation of microscale α-synuclein aggregates called Lewy bodies and Lewy neurites. The morphologies of these structures are typically either neuritic (∼5–10 µm in length) or round (∼5-20 µm diameter),^3^ and have been observed in varying densities in different brain regions depending on the disease stage.^4^ Such structures form the basis of PD diagnostic staging criteria.^4,5^ Further evidence implicating α-synuclein in PD arises from the observation that mutations or gene rearrangements in SNCA,^6–13^ the gene encoding the α-synuclein protein, cause early-onset autosomal dominant PD and variants in the SNCA gene increase the risk of sporadic PD.

Protein aggregation occurs through the self-assembly of monomeric α-synuclein into small protein assemblies, which undergo growth and structural conversion to soluble intermediate species, gradually acquiring cross β-sheet structure.^14–16^ The smaller intermediary structures, referred to as oligomers, are assemblies between monomers and the much larger fibrillar structures found in Lewy bodies.^17^ Typically, oligomers are expected to contain 10s to 100s^18^ of monomeric protein units. Oligomers can have a variety of post-translational modifications and conformations and contain α-helices and/or β-pleated sheets.^19–21^ In cell culture and animal models, it has been shown that oligomers cause neurotoxicity and neuronal death consistent with PD.^22–26^ Oligomers have mainly been studied using recombinant protein, but these aggregates are not identical to the oligomeric species found in human tissue, and different preparation protocols for recombinant oligomers can lead to a variety of characteristics^17,27–29^, motivating studies on native proteins. Detecting endogenous, small aggregates in post-mortem human brain has long remained elusive, primarily due to a lack of sensitivity. Proximity-ligation assays verified the presence of small α-synuclein aggregates by signal amplification,^16^ however, the reporter in this assay does not conserve the native oligomeric structures and may disproportionately amplify larger aggregates.^16,30^ Thus, direct visualisation of oligomers in brain tissue has, so far, not been possible, hindering our understanding of how oligomeric species are distributed spatially and by size.

Here, we present an optical detection and analysis platform, ASA-PD (Advanced Sensing of Aggregates—Parkinson’s Disease), that can be used to quantify aggregate density, distribution, and size directly in fluorescently labelled post-mortem human brain tissue. We applied ASA-PD to characterise α-synuclein single oligomers in large areas of post-mortem tissue sections from patients with PD and matched healthy controls (HC). We detected and characterised over 1.2 million α-synuclein oligomers. To our knowledge, these data represent the first large-scale direct visualisation of α-synuclein oligomers in human brain tissue. The entire dataset, metadata, and analysis toolset have been made available online (see Data Availability). In addition to the microscale aggregates described by classical Lewy pathology,^31^ our data show that vast numbers of oligomers are present in both PD and HC samples. Despite a similar density of oligomers, PD samples contained a population of specific oligomers largely absent from the HCs. This population is preserved across different brain banks, disease stages, immunofluorescent labels, antigen-retrieval methods, and sample-preservation methods, namely fixed and frozen tissue. This finding is consistent with the hypothesis that misfolded α-synuclein oligomers readily form a continuum of larger nanoscale aggregates that eventually give rise to the microscale structures traditionally associated with the disease. We name this population *disease-specific oligomers* and characterise their distribution in neurons and microglia using multicolour fluorescence imaging, revealing further differences in cell type populations.

## Results

### Autofluorescence suppression and high-sensitivity microscopy enable oligomer detection in human brain tissue

An overview of the ASA-PD pipeline is shown in Figure 1. In brief, the aim is to capture spatial data over the entire size scales of structures most critical in PD, from individual cells to small aggregates (Figure 1a). Detailed descriptions of the sample preparation steps are described in the Methods: Immunofluorescence tissue preparation. 8 μm-thick brain tissue sections are mounted on a glass slide, stained and then processed in the five stages illustrated in Figure 1b: (1) background suppression, (2) enhanced imaging, (3) feature detection, (4) analytical computation, and (5) spatial distribution analysis, where the first two steps contain the experimental portion of our workflow, and steps 3-5 perform image-processing tasks and analysis.

**Fig. 1.**
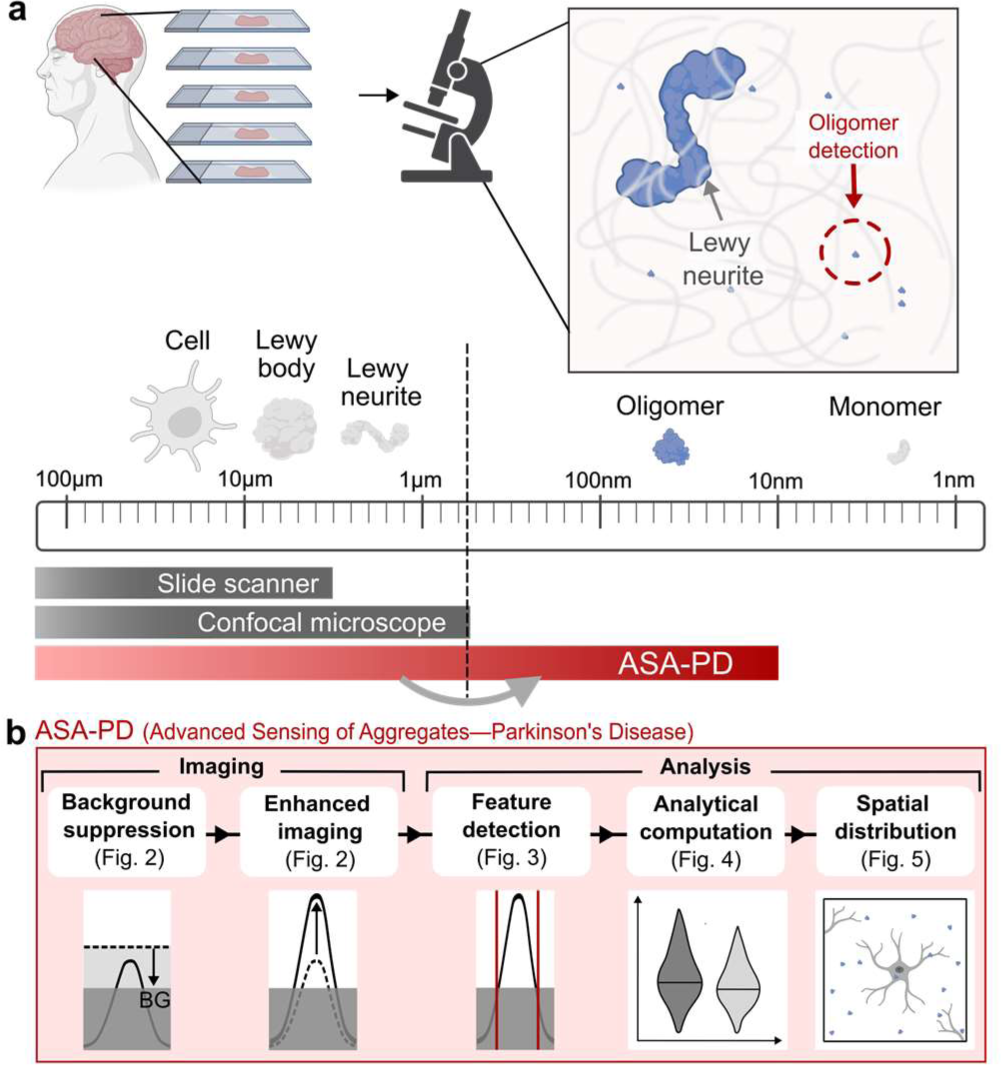
ASA-PD (Advanced Sensing of Aggregates—Parkinson’s Disease). **a**. ASA-PD is an imaging and analysis method for detecting protein aggregates in tissue down to nanoscale oligomers. **b**. The five main steps for imaging and analysis. Background suppression and enhanced imaging improve the signal-to-noise ratio such that aggregates can be detected and quantified in the analysis pipeline including the spatial distributions relative to cell-specific markers. Created in part using biorender.com.

Detecting oligomers and protein aggregates is relatively routine *in vitro* conditions^32–34^ but very challenging in *vivo* due to the poor signal-to-noise ratio in tissue. High background intensities, caused by tissue autofluorescence, act as a noise floor that obscures the presence of dim objects like oligomers. This noise effectively implements a brightness filter, leaving only large protein aggregates, with many attached fluorescent antibodies, as detectable species. To reduce the high autofluorescence of human brain tissue that inhibits sensitive imaging, we deployed Sudan Black (Figure 2a), a fat-soluble diazo dye and well-known autofluorescence quencher on untreated brain tissue sections.^35^ Under optimised conditions, 10 minutes of incubation with 0.1% Sudan Black led to a 93% reduction of background autofluorescence for 561 nm laser excitation (26 W/cm^2^ illumination intensity), corresponding to a decrease in median detected photon counts from 4,400 ± 1040 photons ± median absolute deviation, (MAD) to 333 ± 47 photons (Figure 2b, the background reduction for other excitation colours (488, 561, 641 nm), treatment times and concentrations are shown in Supplementary Figure S1). Next, we repeated this background suppression step on antibody-labelled samples, testing antibodies for α-synuclein and its phosphorylated form (Supplementary Figure S2). The background reduction by Sudan black facilitated the reliable detection of small features in images with a vastly improved signal-to-noise ratio, as shown in Figure 2c.

**Fig. 2.**
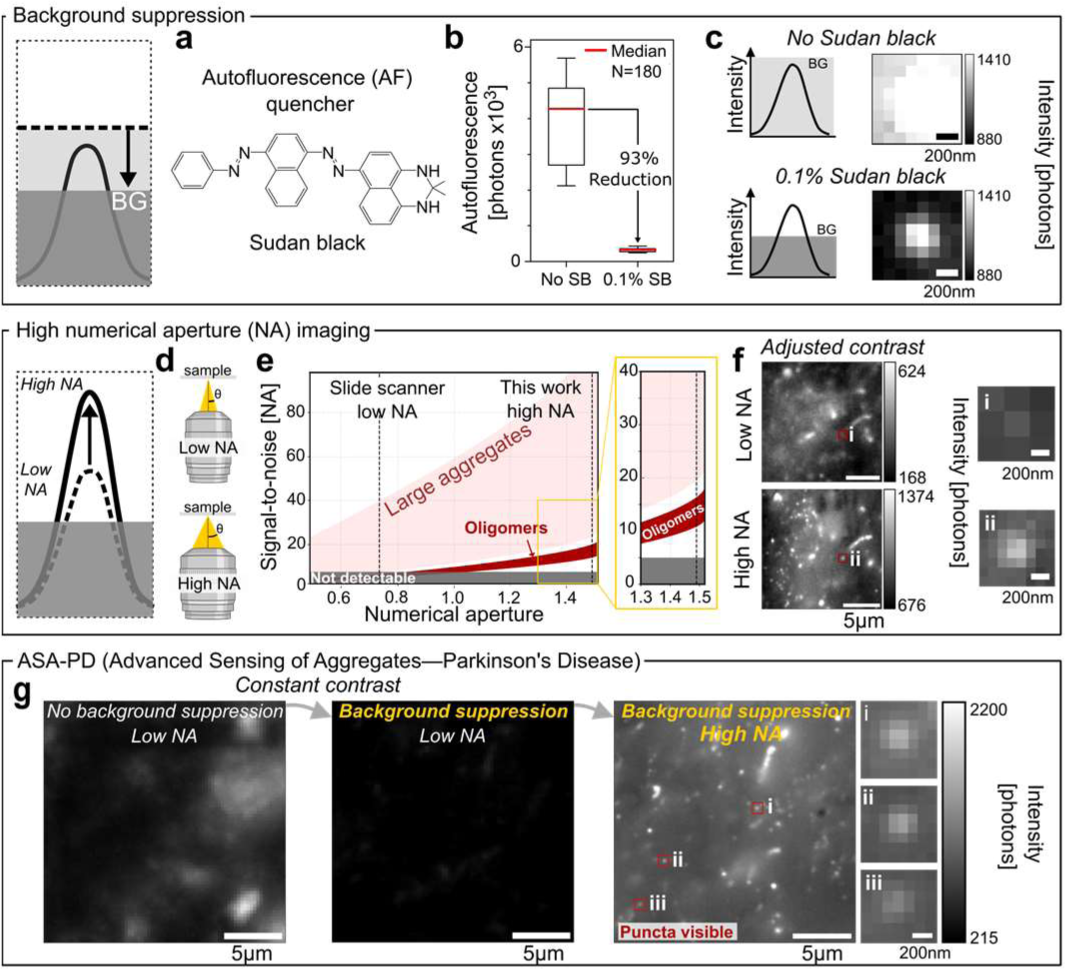
Background suppression and high-sensitivity microscopy in human brain tissue. **a**. Background suppression is achieved with an autofluorescence quencher, Sudan Black (SB). **b**. The median autofluorescence (AF) after treatment with 0.1% SB, N=180 images per sample. **c**. Before quenching, the fluorescence from Alexa fluor 568 labelled small aggregates is masked by the background autofluorescence. After quenching, small aggregates can be easily visualised (both images excited at 561nm, 26 W/cm^2^). **d**. A high numerical aperture (NA) objective collects a larger amount of light from the sample. **e**. Modelled signal-to-noise ratio for imaging punctate objects in post-quenched tissue background at 100*×* magnification across a range of numerical apertures (NA) of objectives. Only at high NA (>1) large aggregates and oligomers become detectable. **f**. Images of p-syn stained PD tissue with 40x mag, NA=0.75 (top) and 100x mag, NA=1.49 (bottom). Closeups shows the same small aggregate is clearly visible at high magnification and high NA. **g**. Images of p-syn stained PD tissue with no background suppression and low NA (left), background suppression implemented and low NA (middle) and background suppression implemented and high NA (right). Several example puncta are shown in the closeups (oligomers) after background suppression is implemented and a high NA objective is used. Created in part using biorender.com.

To visualise the α-synuclein aggregates most associated with PD, we used an antibody targeting phosphorylated α-synuclein at Serine 129 (hereafter called p-syn). This post translational modification promotes inclusion formation and/or toxicity in human cells^36^, Drosophila^37^, and rodent models^38,39^. Further, p-syn forms the vast majority of all insoluble α-synuclein aggregates in the PD brain.^40^ Given this link between p-syn and pathology in synucleinopathies, we tested two complementary antibodies targeting the pS129 epitope of α-synuclein raised in two species (rabbit, AB_2270761, and mouse, AB_2819037, see Methods, Supplementary Tables 3&4). Both antibodies showed characteristic Lewy pathology in both DAB and immunofluorescence staining and were shown to be specific through significant co-localisation with a second antibody for total α-synuclein (AB_2832854). The final p-syn antibody selection, AB_2819037, was based on several criteria: (1) the degree of coincidence of the p-syn antibody with an antibody directed towards total α-synuclein; (2) co-location with other disease-related proteins, such as ubiquitin and p62; and (3) the demonstration of antibody specificity for human α-synuclein based on mouse tissue with overexpression or knockout of human α-synuclein (Supplementary Figures S2).

One way to improve the signal-to-noise ratio beyond reducing the overall background intensity is by improving the light-collection efficiency of the imaging system. In most microscopes, the least efficient step in light collection occurs at the objective lens of the microscope and is encoded in the numerical aperture (NA). Using a high NA objective lens has two main impacts: first, the NA scales with the collection angle of collected light^41^ (Figure 2d, Supplementary Equation S9), and thus, more precious photons from the sample are collected at high NA. Second, increasing the NA improves the image resolution^42^ Supplementary Equation S5). For oligomer imaging in tissue, we deployed a 1.49 NA oil-immersion, 100 × microscope objective lens often used in single-molecule fluorescence applications.^43^ The result is an overall increase in the signal-to-noise ratio for all objects, which is particularly important for the oligomeric species that fall below the detectability range for lower NA objectives (Figure 2e), such as the air objectives most often used in slide scanners for clinical applications.^44^ Figure 2f compares a 0.75 NA 40 × air objective lens (top) with the 1.49 NA 100 × oil objective lens used in this study (bottom) for the same tissue sample stained for phosphorylated α-synuclein and quenched with 0.1% Sudan Black. The effect of background suppression and increased light collection using a high NA are shown in Figure 2g. In these images, a wide variety of object sizes become visible, which we divide into two classes based on their apparent size. We define “**large aggregates”** as objects larger than the optical diffraction limit of visible light, spanning 200nm to 10s of microns, and “**oligomers”** as objects with a physical size below the optical diffraction limit (<200 nm). The fluorescence signal from the latter manifests as a small symmetric puncta in the image. Three example oligomers that become visible via ASA-PD are highlighted in Figure 2g_i-iii_.

Applying the ASA-PD protocol in tissue revealed hundreds of detectable puncta per a typical field of view (55 × 55 µm^2^) in both PD and HCs (Supplementary Figure S3). To perform statistically robust comparisons between samples, we developed a computationally efficient method for detecting and quantifying fluorescent species. This open-source analysis pipeline^45^ facilitates the rapid processing of large image libraries, facilitating transparent, shareable, and verifiable results. A schematic illustrating the analysis method and its validation is shown in Figure 3. In brief, the analysis pipeline identifies features in an image, classifies them as either large aggregates or oligomers, and quantifies details such as brightness, size, and position.^45^ A detailed description of the analysis is provided in Supplementary Information Note 1: Aggregate-detection pipeline and Supplementary Figures S4-S10. Figure 3a shows a typical PD image containing nano and microscale features. Microscale aggregates, such as Lewy bodies and Lewy neurites, are extremely bright in the dataset. These objects can be segmented with a simple intensity threshold after a background subtraction step (see *Large-object pipeline*, Figure 3b, Supplemental Figure S5). As large objects are sometimes extended over multiple z-slices, the masks in each plane were multiplied by a segmented maximum-intensity projection from the z-stack. Smaller aggregates appear as dim, diffraction-limited puncta, and it is crucial to account for local background heterogeneity for detection (see S*mall-aggregate pipeline*, Figure 3b). To do so, a Ricker wavelet filter,^46^ which highlights features of a desired size, was applied to each image. Next, a threshold was used to create a mask containing only small objects. Those objects with a footprint larger than the diffraction limit, as defined by the Rayleigh criterion,^41^ were reclassified as *Large* for subsequent analysis. Finally, the large and small aggregate masks were compared, and overlapping objects were removed from the oligomer dataset. Figure 3c shows an overlay of the detected objects on the original image, and a gallery of diffraction-limited puncta, that is, oligomers, is shown in Figure 3d.

**Fig. 3.**
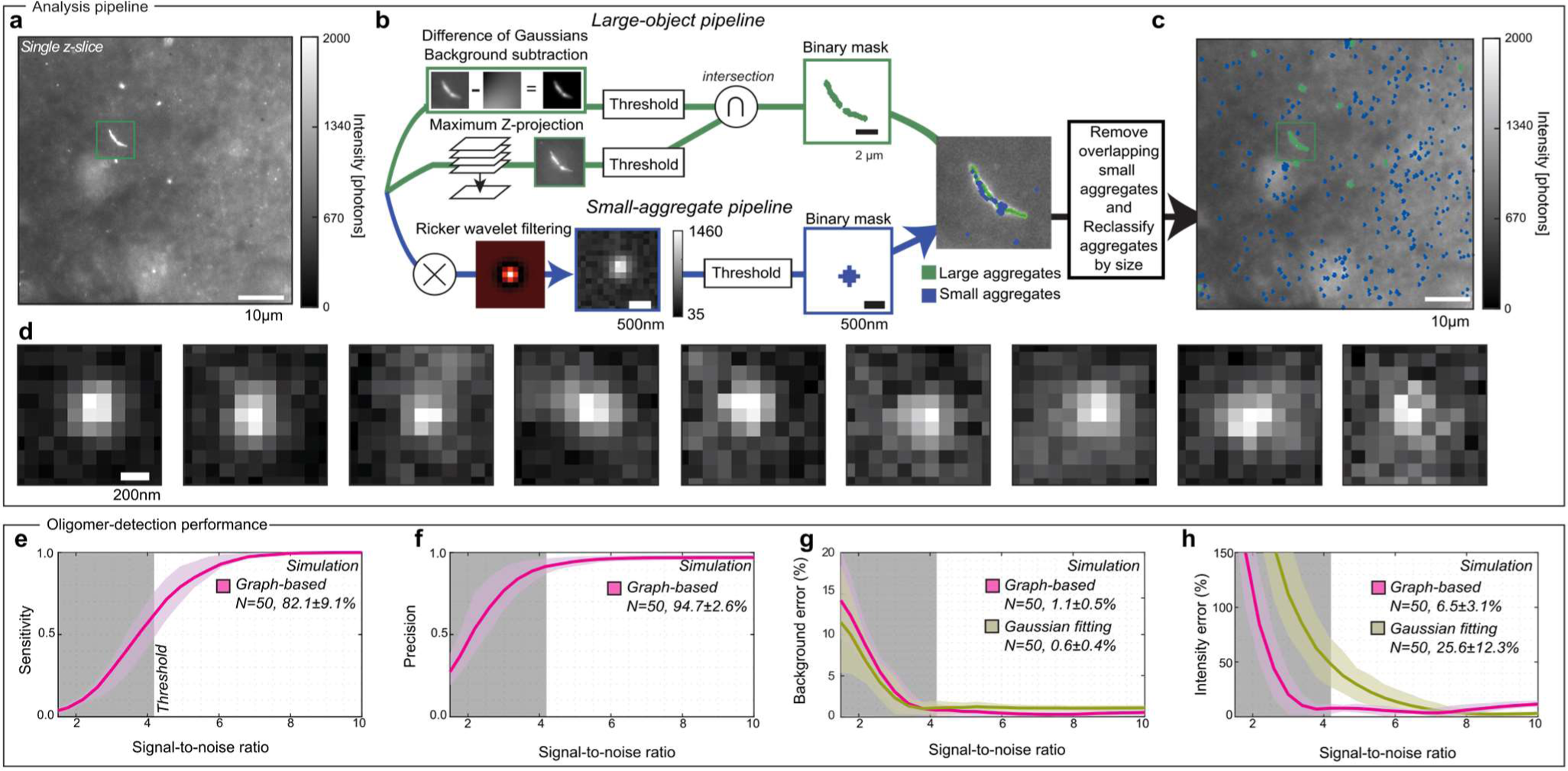
Aggregate-detection pipeline. **a**. A typical sample image containing features of various sizes and intensities, i.e. Lewy neurites, micron-scale aggregates, and sub-diffraction-limit oligomers. **b**. The aggregate-detection pipeline for measuring large aggregates (top) and sub-diffraction-sized features (bottom). The large-object pipeline combines the z-projected intensity data with background-subtracted and threshold individual slices to generate a binary mask. Small aggregates are identified using a Ricker-wavelet filter that acts as a bandpass, emphasising small spots, which are then measured with a threshold and sorted by the number of pixels above the background. Features larger than the diffraction limit are reclassified as “large” and features overlapping between the two masks are removed from the small aggregate pool. **c**. The large (green) and oligomer (blue) masks shown over the original image. **d**. representative oligomers detected from (**c**). **e–h**. Quantification of the pipeline performance using simulated images of diffraction-limited spots on a noisy background at various signal-to-noise ratios. The intensity and average background values for all detected peaks in simulated images were estimated by quantifying the pixel values around the detected peaks (pink curve), and by fitting a symmetric 2D-Gaussian function with nonlinear least squares fitting (brown). The gray box represents the lower threshold bound for analysis.

To evaluate the performance of the pipeline for detecting and characterising oligomers, we simulated images of puncta on noisy backgrounds at various signal-to-noise levels based on empirically determined parameters (Supplementary Note 2: Simulations). In the signal-to-noise range of our data, approximately ∼4-to-6, the algorithm’s sensitivity was >82%, with a precision of >94% (Figure 3ef). At the same time, the relative error for estimating the local background, 1.1%, and brightness, 6.5%, per oligomer outperformed Nonlinear least-squares Gaussian fitting in this low signal-to-noise regime (Figure 3gh), where Gaussian fitting performed poorly on aberrantly detected pixels, i.e., false positives.

### ASA-PD reveals a subpopulation of disease-specific α-synuclein oligomers

To characterise the distributions of α-synuclein in brain tissue, we selected three PD brains (Braak stage 6) and three HC brains for imaging (Supplementary Table 1). In total, 15 PD and 15 HC tissue sections from the anterior cingulate cortical gyrus (ACG) were put through the ASA-PD process. At this point, three principal regions within the grey matter were selected for investigation. At each of these regions, nine fields of view were captured in a 3 × 3 grid, with a lateral separation of 150 μm to avoid any spatial overlap (each image covers 55 × 55µm^2^) in 17 axial planes using a 500 nm step size (Figure 4a, see Methods: Microscopy for microscope-automation details). This process generated 13,770 high-resolution images (>41.6 mm^2^) that were manually validated to ensure the sample was in focus and the tissue contained no significant tears or defects. After this verification step, 12,028 images remained, 87.5% of the original dataset (5,954 PD and 6,074HC images). These images were analysed as described in the previous section to map large aggregates (Figure 4cd) and oligomers (Figure 4ef). Negative control experiments, lacking primary antibodies, were performed in PD tissue to quantify the degree of false positives caused by residual autofluorescence and unbound secondary antibodies (Figure S10).

**Fig. 4.**
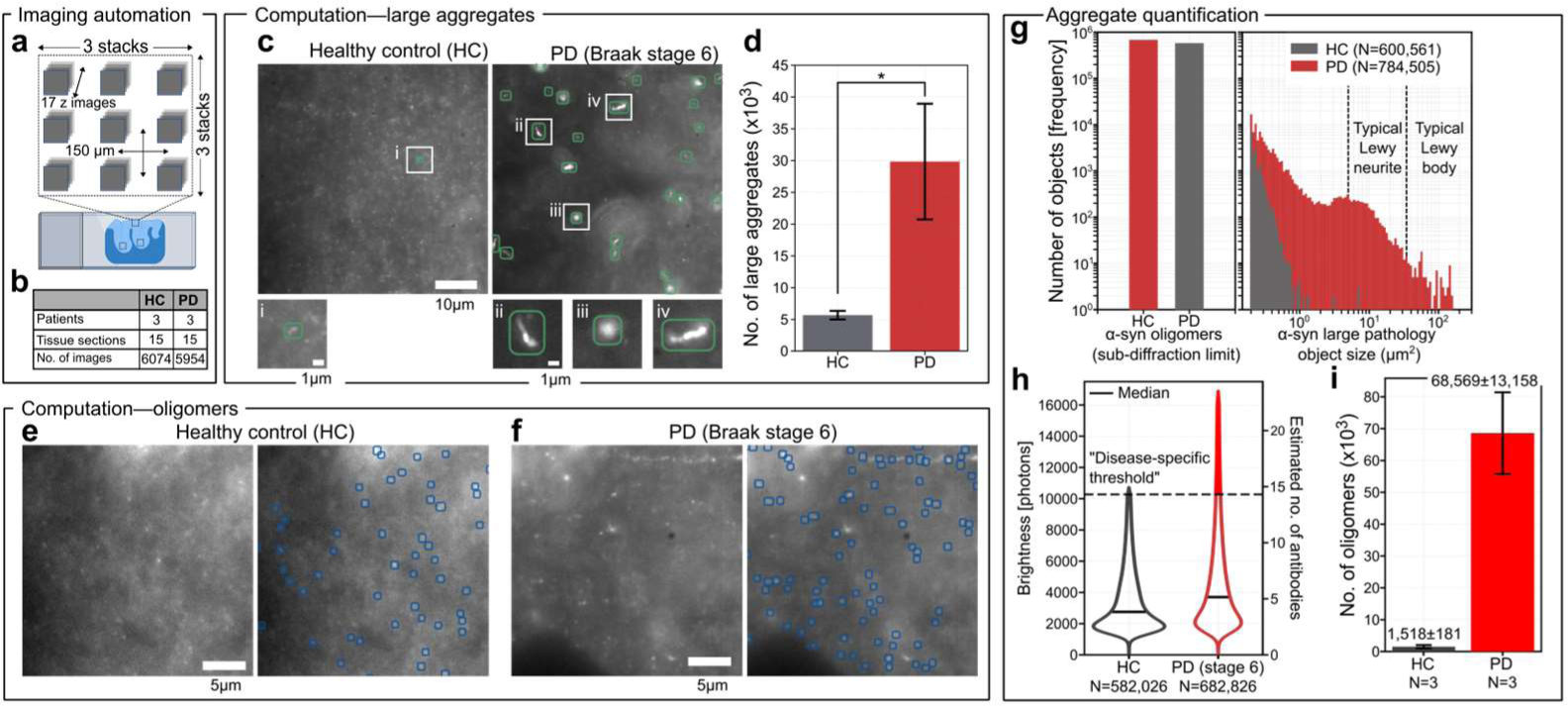
Aggregate distributions in human brain tissue. **a.** Imaging of grey matter was performed in three areas, each area being a 3×3 grid of z stacks (17 slices) spaced by 150 µm. **b**. Number of HC and PD patients (n = 3), number of tissue sections (n = 15), and number of images taken NHC = 6,074 and NPD = 5,954. **c**. Examples of analysed field of views showing only the detected large aggregates. **d.** Number of large aggregates detected per patient over 1800 field of views (5.4 mm^2^). Mean ± S.D. for large aggregates was 1,751 ± 229 in HC and 18, 673 ± 7929 in PD. **e.f.** Example field of views of detected α-synuclein oligomers in HC and PD (Braak stage 6), respectively. **g**. Total number of α-synuclein aggregates in HC and PD tissues. Left panel shows oligomers (< 0.04 μm^2^). Right panel shows large aggregates (> 0.04 μm^2^). The typical Lewy neurites sizes, ∼5 to 30 μm^2^, and Lewy bodies, ∼30 to 300 μm^2^, are shown for reference. **h**. Violin plot of brightness of detected oligomers truncated at 1.5 x IQR. Oligomers in HC had a median of 2,750 photons (MAD=1060) and of 3,700 photons (MAD =1690) for PD. The bright subpopulation of oligomers is shown in red for PD. **i**. The total number of detected oligomers per patient above this brightness threshold, 10,280 photons. Error bars are variation in boundary rejection % per patient, propagated. Created in part using biorender.com.

From the ∼12,000 images recorded across 30 tissue sections, we obtained a dataset containing more than 125,000 large aggregates and 1,260,000 oligomers.^47,48^ From the ∼400 fields of view ∼1.2 mm^2^, from each patient sample, the average number of large aggregates detected was ∼10 × more in PD patients than in the HC, with 18,673 ± 7929 in PD and 1,751 ± 229 in HC, respectively, see Figure 4d. These aggregates were distributed over a broad range of sizes from 0.04-100µm^2^ in PD and 0.04-1µm^2^ in HC (Figure 4g), where the aggregate sizes associated with Lewy pathology were essentially exclusively found in PD samples, consistent with the original tissue classifications (Supplementary Table 1). In the detected oligomer populations, the total number of aggregates in PD and HCs were much more similar (Figure 4g), with 682,826 and 582,026 oligomers, respectively (0.082 oligomers/μm^2^ for PD and 0.067 oligomers/μm^2^ for HC).

Below the diffraction limit, it is not possible to measure the aggregate size directly; however, for larger aggregates, where the size and brightness can be measured, the two were strongly linearly proportional (Supplemental Figure S11). We, therefore, characterised the distribution of oligomer intensities as a proxy for their approximate size (Figure 4h). This number was converted to the approximate number of bound secondary antibodies, as each antibody contributes ∼700 photons in our imaging conditions (Supplementary Figure S12). The aggregate brightness distributions for all measured objects are shown in Supplementary Figure S11. Relative to the HC, the median was larger for PD samples, 3,700 photons (MAD =1690) and 2,750 photons (MAD =1060), and the distribution of brightnesses in PD samples was also broader, characterised by its interquartile range IQR_PD_ = 4,280 photons compared to IQR_HC_ = 2,690 for HCs. To determine if the distribution tail was reproducibly different between PD and HC samples, we defined and applied a brightness threshold that separated the disease-specific population Figure 4h. The number of oligomers above this threshold (10,280 photons, ∼15 antibodies) is shown in Figure 4i. This data represents ∼10% of all measured PD oligomers but only 0.26% of all HC oligomers (68,569 in PD and 1,518 in HC). The existence of this bright, disease-specific subpopulation was highly robust and was observed consistently using different α-synuclein antibodies, measured in different disease stages, from different brain banks, across multiple individuals, and using different antigen retrieval methods (Supplementary Figure S13, Supplementary Tables 1&2).

### Oligomer density varies across cell types

In addition to measuring the oligomer densities and size distributions, ASA-PD can be used to analyse spatial distributions and overlapping signals in co-stained samples (Figure 5). This capability is important because the large α-synuclein inclusions have been found predominantly in neurons in Parkinson’s disease.^3,4,49^ To validate if our pipeline could detect differences in the α-synuclein density for particular cell types, we co-stained samples with α-synuclein and cell markers such that fluorescence from cell marker, where the signal-to-noise is relatively high, did not interfere with the high-sensitivity imaging in the oligomer channel, *i.e.,* we minimised channel crosstalk. Specifically, neurons were labelled by staining neurofilaments (NF) with Alexa Fluor 488-conjugated antibodies, while phosphorylated α-synuclein was labelled with Alexa Fluor 568 dyes. Images were recorded separately with excitation intensities of 2.4 W/cm^2^ 488 nm and 25.9 W/cm^2^ 561 nm. The 488 nm excitation intensity was kept low to prevent photobleaching of the oligomers during the z-scan, as the high density of cell markers was sufficient to resolve the structures (Figure 5ad).

**Fig. 5.**
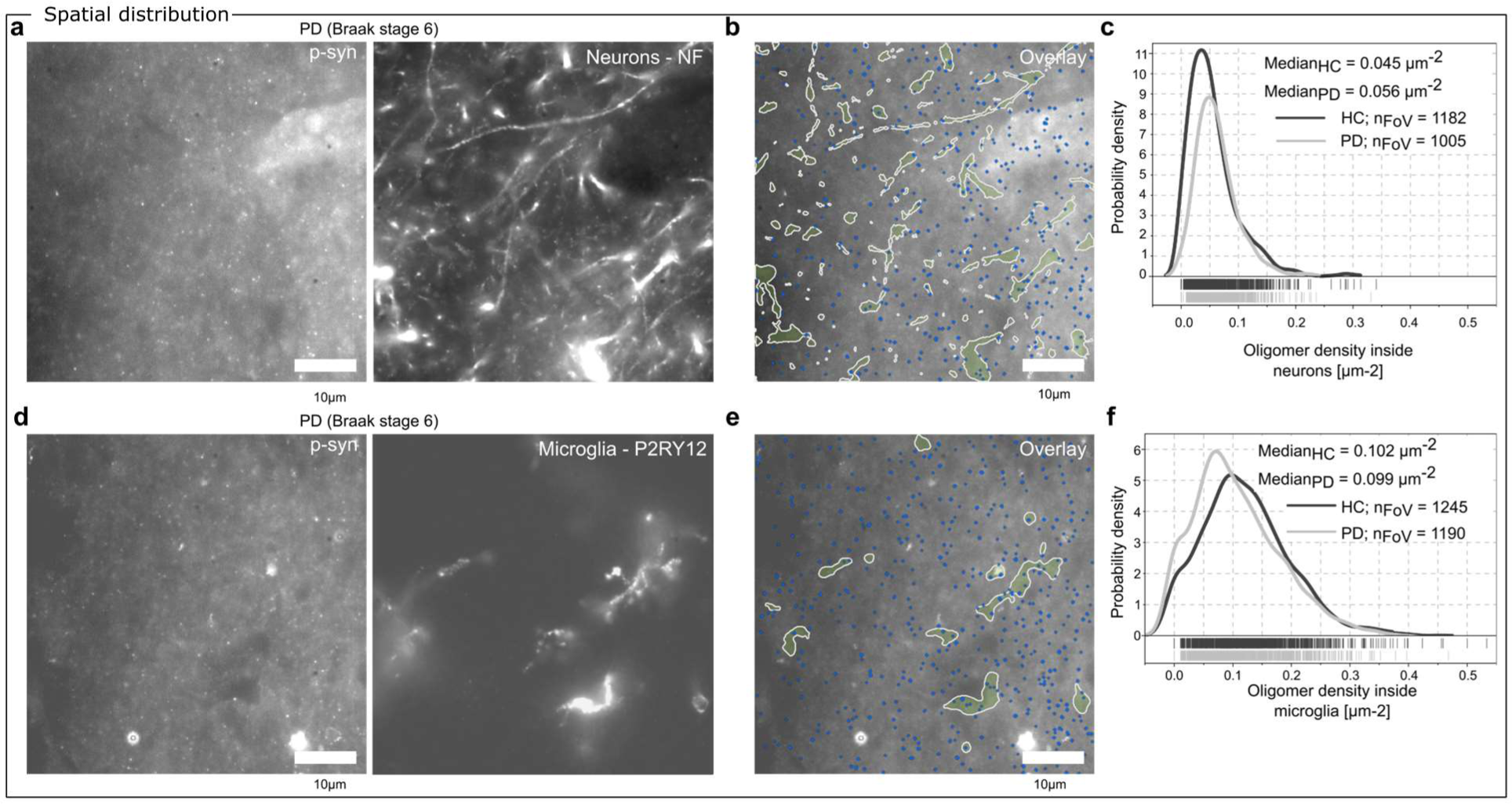
Spatial distribution of oligomers with respect to different brain cell types. **a**. Example field of view of co-stained tissue from PD (Braak Stage 6), α-synuclein (p-syn) (left) and neurons, (neurofilaments, NF) (right). **b**. Overlay of detected oligomers (blue) and detected neuron boundaries (yellow). **c**. Oligomer densities inside neurons for 3 PD patients (1182 field of views) and 3 HC (1005 field of views) with median density of 0.0451 µm^−2^ (MAD = 0.023 µm^−2^) in HC and 0.0563 µm^−2^ (MAD = 0.021 µm^−2^) in PD. **d**. Example field of view of co-stained tissue from PD tissue (Braak Stage 6) with α-synuclein (p-syn) (left) and microglia (P2RY12) (right). **e**. Overlay of detected oligomers (blue) and detected microglia boundaries (yellow). **f**. Oligomer density analysis of oligomers inside microglia across 3 PD patients (1245 field of views) and 3 HC (1190 field of views) with median density of 0.102 µm^−2^ (MAD = 0.046 µm^−2^) in HC and 0.099 µm^−2^ (MAD = 0.046 µm^−2^) in PD.

Three PD patients (Braak stage 6) and three HCs were stained for neurons (neurofilament, NF) and co-stained for p-syn, see Figure 5a. By implementing oligomer detection and cell segmentation in our imaging pipeline (Figure 5b), we were able to analyse the density of oligomers inside NF-positive neurons for 2,187 fields of view. We found that the density of oligomers inside neurons increased in PD to a median density of oligomers of 0.0563 µm^−2^ (MAD = 0.021 µm^−2^) when compared to HCs with a median density of oligomers of 0.0451 µm^−2^ (MAD = 0.023 µm^−2^), see Figure 5c.

Continuing this approach into an alternative cell type, we also stained for a different cell type in the same region of brain tissue. Here, we immunolabelled P2RY12, a microglia marker (excited at 488 nm), alongside p-syn (excited at 561 nm), see Figure 5d. Across 6 tissue sections from individual patients, 3 PD and 3 HC, 2,435 fields of view were imaged. We found that there was a slight decrease in the median density of oligomers inside microglia in PD, 0.099 µm^−2^ (MAD = 0.046 µm^−2^) when compared to HC 0.102 µm^−2^ (MAD = 0.046 µm^−2^). We have demonstrated that ASA-PD makes it possible to determine the oligomer density inside and outside cells. However, more samples will be required in the future to conclude the cell specificity of oligomer formation. Nonetheless, these data demonstrate that ASA-PD can be used in future studies to uncover the spatial organisation of oligomers in cell types in the PD brain.

## Discussion

We have demonstrated the direct detection of single α-synuclein oligomers in human post-mortem brain tissue and performed quantitative analysis of more than 1.2 million oligomers across 30 tissue sections (682,826 oligomers detected across 15 PD tissue sections, whereas compared to 582,026 oligomers detected across 15 HC sections). The acquisition of this large-scale dataset was made possible by the imaging component of ASA-PD, which is a combination of background suppression and high-NA collection of light that improves the signal-to-noise sufficiently to visualise the dim signal from individual nanoscale aggregates. The analysis pipeline that forms the detection step of ASA-PD allowed sensitive and precise detection of these dim signals, as well as the ability to determine cell-specificity from co-staining and cell segmentation. The detection of small protein assemblies in post-mortem tissue is sensitive to many experimental fluctuations, and therefore ASA-PD was tested in a range of conditions, including the source of the brains (from two independent brain banks, Supplementary Table 2), the sample preparation (formalin-fixed paraffin-embedded (FFPE) samples *vs.* frozen samples), and the traditional antigen retrieval methods (formic acid and heat mediated epitope retrieval). We found that the most consistent detection of oligomers was obtained using FFPE Braak stage 6 tissue sections from a single brain bank, without any formic acid antigen retrieval; however, broadly similar trends were observed in all conditions tested.

In agreement with the classical studies of neuropathology in Parkinson’s disease post-mortem brain, we observed a six-times increase in the number of large aggregates in PD with respect to HC samples. Lewy pathology, consisting of Lewy bodies and Lewy neurites, are microscale aggregates that are always found in sporadic PD cases (and have been used to define the disease^4,50^) and are sometimes found in HC tissue, where it has been described as incidental Lewy Body Disease.^51^ Such studies have described the presence or absence of large Lewy aggregates, reporting an estimated level of abundance as a marker of severity of Lewy pathology which lacks detailed quantitation. The technical advances implemented in ASA-PD have allowed us to capture the entire size range of protein aggregates containing phosphorylated α-synuclein and the quantification of their sizes and frequency in human Parkinson’s disease tissue.

With this large-scale dataset, we established that there is a continuum of aggregate sizes in the human brain, ranging from large (microscale) to very small (nanoscale). Using ASA-PD, each stage of the protein aggregation pathway present in disease brain can now be measured. Importantly, large aggregates are much less frequent compared to small aggregates, as there is 16-times increase in small aggregates below the diffraction limit of light compared to larger ones in disease. Therefore small aggregates are by far the more abundant aggregate species, prompting the question about the nature of their link with PD, a longstanding unanswered question in the field. Small aggregates, or oligomers, have been previously shown to be increased in brains with Lewy pathology compared to controls,^52–54^ and elevated levels of oligomers have been detected in the cerebrospinal fluid of PD patients compared to controls.^55–58^ Furthermore, a PLA-based approach revealed a widespread distribution of α-synuclein oligomers in disease brain tissue, in contrast to a highly restricted distribution of Lewy-related pathology.^59^

ASA-PD revealed an overall abundance of oligomers/small aggregates in both control and Parkinson’s disease tissue (equating to hundreds of oligomers/cell). Their presence in control and disease suggests that small α-synuclein aggregates oligomerise under physiological conditions, where their formation and clearance are kept in balance by the protein homeostasis system. For instance, the existence of α-synuclein as a tetramer or higher-order conformations has been previously reported in healthy-control samples using biochemical approaches.^60^ Furthermore, serine 129 phosphorylation (pS129) may arise under homeostatic conditions in response to synapse activity, being a reversible event that may regulate neuronal activity.^61^ Activity-induced pS129 leads to conformational changes that facilitate interactions with new binding partners at the synapse, enabling α-synuclein to attenuate neurotransmission throughout regulating neurotransmitter release.^62^.The abundance of small assemblies, or oligomers, of phosphorylated α-synuclein in both healthy and PD brains would be in keeping with these proposed physiological roles for pS129 α-synuclein.

Although α-synuclein oligomers are present in similar numbers both in PD and HC, because of the large oligomer statistics accessible through ASA-PD, we have been able to identify a clear difference - a subpopulation of 9.7% - of the total oligomers that are larger and found only in PD brains. Therefore, these results suggest that a small proportion of the physiological oligomers that are detected in HC may undergo a transition to the pathological large oligomers detected in PD. Once this transition has occurred, these pathological oligomers may then ultimately aggregate further and become the fibrillar structures that are found in Lewy bodies and Lewy neurites. This hypothesis is consistent with previous findings^32,63–65^ of aggregation kinetics in vitro, where a small proportion of the total α-synuclein population, under pro-aggregation conditions, will give rise to disease-specific toxic oligomers. In vitro and in human neurons, the earliest oligomers formed are Proteinase K sensitive, relatively inert, and non-toxic (termed Type A), and during aggregation, they undergo a transition to larger, Proteinase K resistant, highly toxic oligomers^15,33,66,67^ (Type B oligomers). This transition is associated with structural conversion from relatively disordered assemblies to highly ordered oligomers with the acquisition of β-sheet structure. It is the acquisition of this β-sheet structure that is associated with the propensity for these in vitro formed pathological oligomers to disrupt membranes and induce toxicity in human neuronal systems.^68^ It is not yet clear whether or how a structural conversion to the disease-specific oligomer, or pathological oligomer, may occur in the brain. However, ASA-PD’s ability to detect a disease-specific oligomer sub-population opens the door to further investigation of their structural properties, specifically their order, sensitivity to degradation, and β-sheet structure.^69^

Using spatial point statistics, ASA-PD did not demonstrate obvious large-scale clustering of the entire oligomer population, consistent with a widespread cellular distribution throughout the anterior cingulate cortex. However, we have shown a small preference for increased oligomeric density in some neurons. As large (Lewy) aggregates are always located within neurons, it is possible that oligomers transition from their control/physiological state into the pathological state (either by size or by structural conversion), where they may precede the formation of the later stage Lewy bodies. How the disease-specific oligomer ultimately forms a Lewy body, containing not only fibrillar α-synuclein but also neuronal lipid membranes and organelles,^70,71^ is not known *in vivo*. Detailed cellular and subcellular imaging of the aggregates species as they transition from the bright disease-specific oligomers to fibrillar structures will provide information on the trajectory of Lewy body formation within neurons in the human brain, their heterogeneity, and the components within them.

This study represents to our knowledge the largest study of its kind to date, involving the analysis of over 1 million oligomers. However, a number of limitations still pose further challenges:

i. *Determination of aggregate sizes*: we have not directly measured the physical dimensions of the oligomers, but inferred the distribution of sizes based on their relative integrated brightnesses. Above the diffraction limit (around ∼200 nm), we observed a strong linear relationship between the integrated brightness of aggregates and their measured area (R^2^ = 0.987, as depicted in Supplemental Figure S11). We hypothesise that this pattern persists below the diffraction limit as well, i.e. to the oligomeric fractions. The linear correlation with area suggests that microscale aggregates have numerous exposed epitopes on their surfaces, making it unlikely for antibodies to penetrate deeply inside the aggregates.
ii. *Detection of distinct oligomer classes*: we validated the use of multiple antibodies, and primarily used an antibody to the Serine 129 C-terminal phosphorylated form of α-synuclein. Therefore, we cannot be certain that the distributions and densities of this specific oligomer class is conserved for other oligomer types. Indeed, antibodies to multiple epitopes, including those targeting the N terminus, have been shown to provide more detailed information on other oligomer subtypes and locations in the brain.^72,73^
iii. *Detailed characterisation of the disease-specific oligomer population*: this will require super-resolution methods or electron microscopy to visualise nanoscale structures. To use super-resolution methods, further signal-to-noise ratio improvement is needed. This could include background reduction, e.g. optical-clearing techniques,^74^ combined with better sectioning, e.g. confocal microscopy or thinner tissue slicing.
iv. *Extended coverage of brain regions*: Higher throughput methods are also needed to increase statistical confidence in biological findings at the macroscopic scale of the brain. While high-throughput methods with single-molecule sensitivity are possible, there are currently no commercially available solutions, necessitating the use of custom-made instruments. We anticipate that approximately a two-order-of-magnitude increase in throughput could enable the exploration of different brain regions, encompass more cases, and facilitate large-scale automation, ultimately establishing a foundational oligomeric map of the Parkinson’s disease (PD) brain.

## Conclusions and Outlook

We designed ASA-PD, a platform technology for the large-scale imaging of protein aggregates in brain tissue. We then illustrated its application to generate the largest dataset to date describing the distribution of α-synuclein aggregates, their prevalence, and their spatial location in the PD brain. Without this quantitative information, it has been challenging to establish the nature of the link between α-synuclein aggregation and Parkinson’s disease. Although substantial evidence from model systems has implicated α-synuclein oligomers in pathological processes, it has remained unclear whether such processes are actually relevant in the disease. With ASA-PD, we identified a small population of specific oligomers that are characteristically present in disease. The ASA-PD platform can thus be used to design mechanistic studies to understand how these disease-specific oligomers are created. These studies will build on the ability of ASA-PD to address the regional and cellular microenvironments that promote the development of disease-specific oligomers, as well as the temporal evolution of the end-stage Lewy body pathology from the disease-specific oligomers. Furthermore, integrating the ASA-PD dataset with other single-cell and spatial RNA and protein technologies will allow the identification of the key pathways and mechanisms associated with the cellular environments that promote oligomer and Lewy body formation. We also note that the ASA-PD method is widely applicable to other neurodegenerative diseases, where the role of protein aggregation remains largely unresolved.

## Materials and Methods

*Detailed protocols are available online*.

-Single-molecule slides for fluorescence microscopy^75^

-Free-floating Mouse brain immunohistochemistry^76^

-Preparing tissue staining and oligomer imaging is available online^77^

-Feature detection software^78^

### Tissue selection

Post-mortem brain tissue was obtained from Queen Square Brain Bank for Neurological Disorders, University College London (QSBB), and Multiple Sclerosis and Parkinson’s Brain Bank, Imperial College London. Braak stage 3/4 PD cases (n=3) were obtained from Imperial Brain Bank, Braak stage 6 PD cases (n=4, 1 for technical controls, 3 for main study) from QSBB and 3 healthy controls from each brain bank to control for brain bank processing effects (n=6, 3 from Imperial, 3 from QSBB) as highlighted in Table 1. Standard diagnostic criteria were used to determine the pathological diagnosis. Case demographics for each case are described in Supplementary Table 1.

### Immunofluorescence tissue preparation

8µm thick formalin-fixed paraffin-embedded (FFPE) and fresh frozen tissue sections from the cingulate cortex were cut from the cases summarised in Table 1 and described in detail in Supplementary Table 1. These sections were loaded onto Superfrost Plus microscope slides. FFPE sections were baked at 37°C for 24 hours and then 60°C overnight. They were deparaffinised in xylene and rehydrated using graded alcohols. Frozen sections were fixed in 4% paraformaldehyde for 30 minutes and washed 3x for 5 minutes in PBS. Endogenous peroxidase activity was blocked in 0.3% H_2_O_2_ in methanol for 10 minutes. All sections underwent heat-mediated epitope retrieval for 10 minutes in citrate buffer (pH 6.0), and half of the sections were additionally incubated in formic acid for 10 minutes prior to heat-mediated epitope retrieval to test which were the optimal antigen retrieval conditions. Non-specific binding was blocked with 10% dried milk solution in PBS. Tissue sections were incubated with primary antibodies; anti-α-synuclein (LB509, AB_2832854 1:100; phospho S129 rabbit polyclonal, AB_2270761, 1:200; phospho S129 mouse monoclonal, AB_2819037, 1:500); anti-P2RY12 (AB_2669027, 1:100); anti-neurofilament (RT-97, AB_2941917, 1:200); anti-Glial fibrillary acidic protein (GFAP) (5C10, AB_2747779, 1:1000); anti-Olig2 (AB_570666 1:100) for 1 h at RT, washed three times for five minutes in PBS followed by the corresponding AlexaFluor (anti-mouse 488, AB_2534069/ anti-mouse 568, AB_144696/ anti-rabbit 488, AB_143165/ anti-rabbit 568, AB_143157 all at 1:200) for 1 hour at room temperature. Sections were kept in the dark from this point onwards. Sections were then washed three times for 5 minutes in PBS and incubated in Sudan Black (multiple concentrations and incubation times were tested, as described in the background suppression section). Sudan Black was removed with three washes of 30% ethanol before they were mounted with Vectashield PLUS (Vector Laboratories, H-1900), coverslipped (22×50 mm #1, VWR, 631-0137), and sealed with CoverGrip™ sealant (Biotium, 23005) for imaging. Sections were stored at 4°C until imaging.

### Single-molecule secondary antibodies

Coverslips (24 x 50 mm, #1, VWR, 48404-453) were argon plasma cleaned (Ar plasma cleaner, PDC-002, Harrick Plasma) for 30 minutes before a trimmed gasket was placed on top (CultureWell™ Reusable Gasket, 6mm diameter, Grace Bio-Labs, 103280). Poly-L-Lysine (0.01 % w/v PLL, Sigma-Aldrich, P4707) was placed in the wells for 30 minutes. The PLL was removed, the wells washed three times with PBS (pH 7.4, 1x Gibco, Thermo Fisher Scientific, 10010023) and the secondary antibody of choice was added (Alexa Fluor 568 goat anti-mouse – AB_144696 or Alexa Fluor 568 goat anti-rabbit – AB_143157) at a dilution of 1:10,000 in PBS from 2 mg/ml stock to a final concentration of 0.2 μg/ml. The antibodies were left in the wells for 5-10 seconds for sufficient surface density before the wells were washed three times with PBS. PBS (30μl) was left in the wells for imaging. Images were taken over two slides with 25 fields of view per slide.

### Microscopy

Images of human post-mortem brain and single-molecule antibodies were taken using a custom-built widefield fluorescence microscope that has been described previously.^79^ Illumination of the sample was by a 488 nm laser (iBeam-SMART, Toptica) and a 561 nm laser (LaserBoxx, DPSS, Oxxius), both of which had the same excitation alignment. Both lasers were circularly polarised using quarter-wave plates, collimated, and expanded to minimise field variation. The laser lines were aligned and focused on the back focal plane of the objective lens (100x Plan Apo TIRF, NA 1.49 oil-immersion, Nikon) to allow for the sample to be illuminated by a highly inclined and laminated optical sheet (HILO). Emitted fluorescence was collected by the objective lens before passing through a dichroic mirror (Di01-R405/488/561/635, Semrock). The collected fluorescence then passed through emission filters dependent on the excitation wavelength (FF01-520/44-25 + BLP01-488R for 488 nm excitation, LP02-568RS-25 + FF01-587/35-25 for 561 nm excitation, Semrock). The fluorescence was then expanded (1.5x) during projection onto an electron-multiplying charge-coupled device (EMCCD, Evolve 512 Delta, Photometrics). The EMCCD was operating in frame transfer mode with an electron multiplication gain of 250 ADU/photon.

Z-stacks of images were taken through the samples in 0.5 µm steps with 17 steps per field of view (FoV) covering a depth of 8 µm. To reduce bias in FoV selection, nine FoVs were recorded in a grid formation of 3 FoVs x 3 FoVs with 150 µm spacing, implemented with the ImageJ^80^ Micromanager plugin.^81^ Three 3 x 3 grids were recorded at three random locations within the grey matter of each section, resulting in 27 z-stacks per section. Images were recorded with 1 s exposure time of the EMCCD. Images were recorded with a power density of 2.4 W/cm^2^ (488 nm excitation) and 25.9 W/cm^2^ (561 nm excitation).

### Camera calibration

To convert the pixel values from counts in analogue-to-digital units (ADU) to photons, a series of images were recorded at different illumination intensities, including one taken under no light to measure the camera (EMCCD, Photometrics, Evolve 512 Delta) offset.^82^ Each intensity was captured in 500 frames, resulting in a total of 3000 frames across the six different illumination levels. For every pixel, the mean and variance were calculated across the 500 frames, generating six different variance and mean values corresponding to the six illumination intensities. The dark counts, i.e. the camera offset per pixel, were determined as the mean pixel value in the dark frame. The camera gain per pixel, expressed in photoelectrons per count, was determined by calculating the slope between the six variance and mean values per pixel, subtracting the dark frame offset.

### Feature detection

Custom code was written to detect both cells and α-synuclein aggregates. Firstly, each z-stack was manually inspected to remove out-of-focus images. For the aggregate channel, two binary masks representing nanoscale and larger fluorescent objects images respectively were created for each image using the following four steps: (1) The positions of large objects were detected with a difference-of-Gaussian kernel (σ_1_ = 2_px_, σ_2_ = 40_px_) and Otsu’s threshold; (2) cellular autofluorescence was removed using high-frequency filtering (σ_high-pass_ = 5_px_), and diffraction-limited-sized features were enhanced through Ricker wavelet filtering (σ_ricker_ = 1.1_px_); (3) bright spots within the image were identified using an intensity threshold based on the top 2.5th percentile of the image; and (4) spurious pixels were removed from the union of the two binary masks in step (1) and (3) *via* morphological opening operation with a disk shape (radius = 1_px_). The resulting mask is then separated into nanoscale and non-nanoscale fluorescent objects, where the threshold was 19_px_, determined by 40nm sub-diffraction limited fluorescent beads (40 nm, FluoSphere F10720). For the cell channel, cell segmentation was achieved using the function described in the first step of the aggregate-detection pipeline but using different kernel sizes (σ) dependent on cell type. For specific details of σs used, see Supplementary Information Note 1. All imaging data was viewed in ImageJ and analysed and plotted using either custom MATLAB code and Origin or custom-written Python code.

## Acknowledgements

This research was funded in part by Aligning Science Across Parkinson’s [ASAP-000509 (MR, NW, MV, SG, SFL), ASAP-000478 (ZJ, JH), ASAP-020370 (PJM)] through the Michael J. Fox Foundation for Parkinson’s Research (MJFF). RA was supported by an Astra Zeneca Postdoctoral Fellowship. CL and PJM was supported by the Medical Research Council (award MC_UU_00003/5), LEW acknolwedges the Canada First Research Excellence Fund (TransMedTech Institute) and the Polytechnique Montréal Direction de la recherche et de l’innovation (DRI) for research support to AD, KB, and LEW. S.G. was supported by Wellcome (100172/Z/12/2) and is currently an MRC Senior Clinical Fellow (MR/T008199/1)

## Author Contributions

Sample preparation: RA, CET, JL, RT, L-MN, CL, ZJ, JH, TL, SG;

Material preparation: RA, CET, JCB, JL, RT, L-MN, CL, AD, SFL;

Data collection: RA, BF, JCB, RT, L-MN CL;

Data analysis: RA, BF, JCB, JSB, GJC, CL, AD, KB, LEW, SG, SFL;

Data interpretation: RA, BF, CET, JCB, JL, RT, JSB, AD, KB, PJM, OJF, BJMT, TL, MR, MV, NWW, LEW, SG, SFL;

Figure preparation: RA, BF, CET, JCB, JL, RT, JSB, LEW, SG, SFL;

Manuscript writing: RA, BF, CET, JL, JSB, MV, LEW, SG, SFL;

Study conception: RA, CET, PJM, TL, MV, NWW, SG, SFL;

Overseeing project: RA, PJM, MV, LEW, SG, SFL;

All authors read and approved the final manuscript.

## Data Availability

All unprocessed data^47^, processed data (binary masks, density and intensity information)^48^, methods, and detailed protocols are available online.

## Declarations

Ethical approval for the study was granted from the Local Research Ethics committee of the National Hospital for Neurology and Neurosurgery.

## Conflict of interest

OJF and BJMT were/are employed by AstraZeneca, OJF is currently employed by MSD. All other authors declare that they have no conflict of interest.

## Supplementary Information

### 1. Supplementary Figures

**Supplementary Fig. S1.**
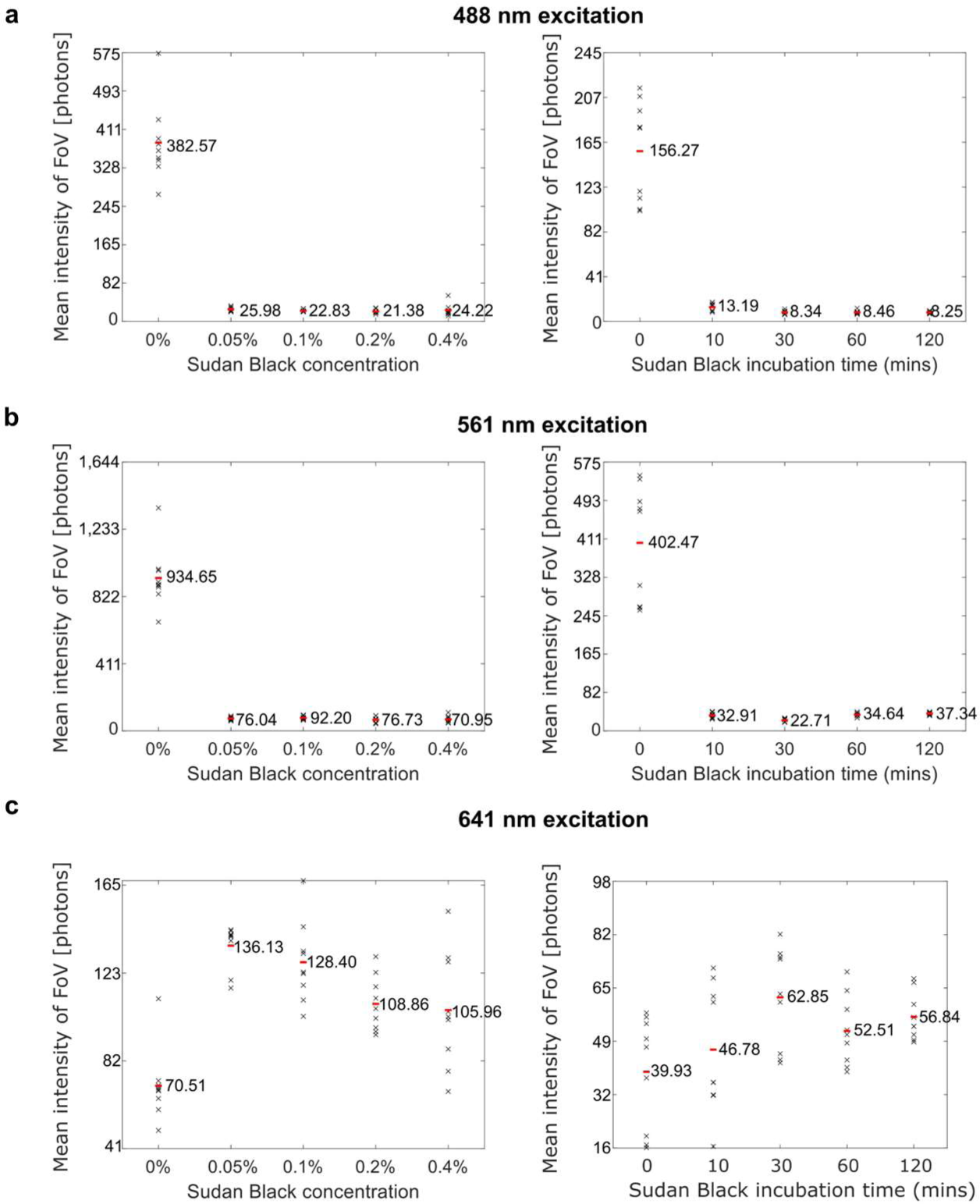
Sudan Black (SB) concentration study and incubation time study. **a.** Data taken at 488 nm excitation, 100 ms, 2.4 W/cm^2^. **b.** Data taken at 561 nm excitation, 1 s, 25.9 W/cm^2^. **c.** Data taken at 641 nm excitation, 100 ms, 7.0 W/cm^2^. Black ‘X’ are an average of 20 FoVs, in 3 different places of grey matter, over 3 sections each from a different Braak Stage 6 PD patient (9 data points per condition). Red is the mean of all ‘X’ for conditions labelled with mean intensity (photons).

**Supplementary Fig. S2.**
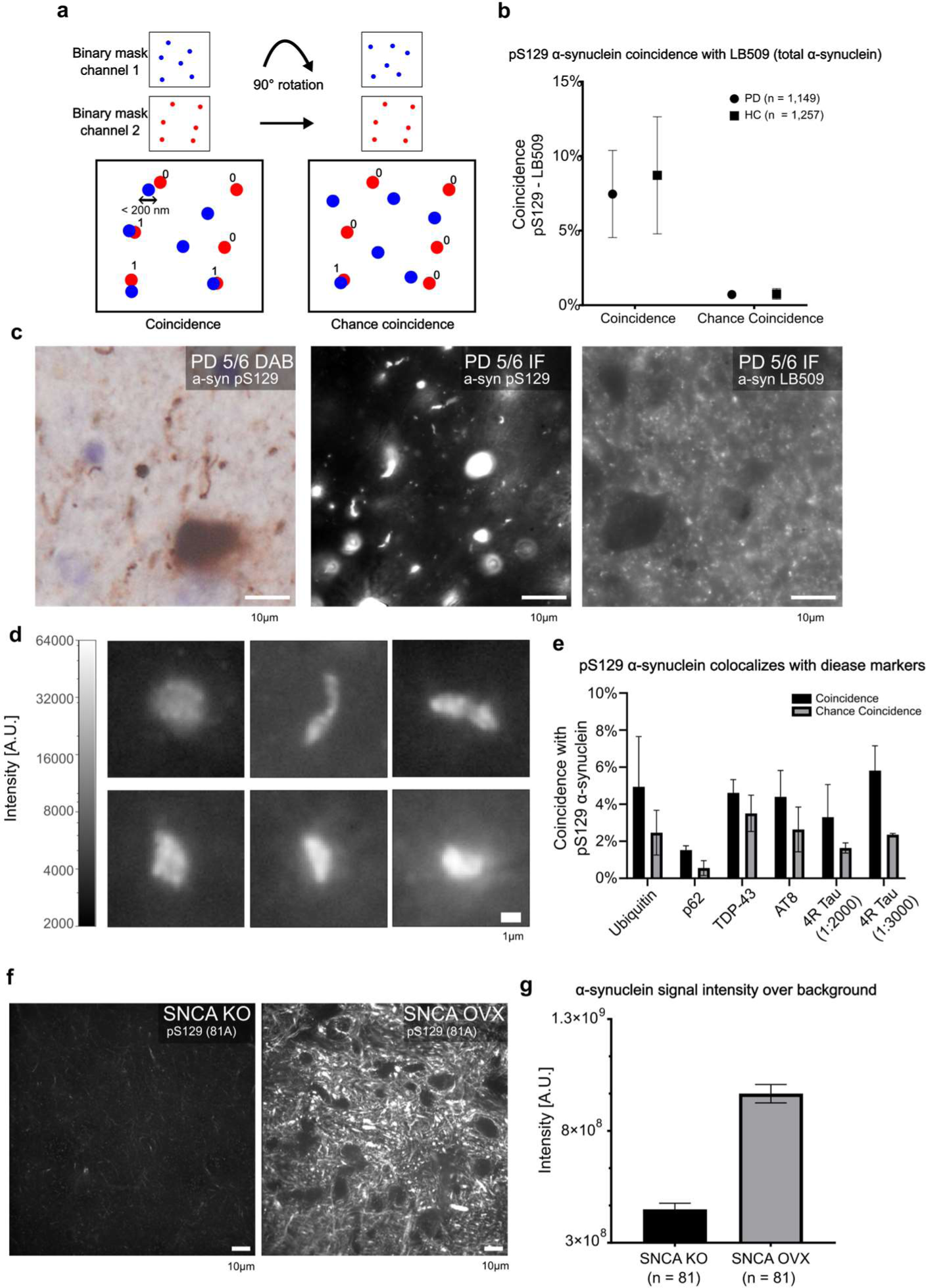
Robust α-synuclein detection with different antibodies. **a.** Schematic representation of the assertion of coincidence of two image channels for diffraction-limited data. Coincidence between both channels is asserted based on an overlay of two binary masks. If *n* > 1 pixel overlaps, a coincidence is registered. Chance coincidence is calculated in the same way after rotating one binary mask 90 degrees. **b.** Coincidence and chance coincidence of LB509 (AB_2832854) on pS129 α-synuclein (AB_2270761) across n = 3 Parkinson’s disease Braak stage 5/6 and n = 3 HCs. The *n* in the legend represents the number of fields of view. Error bars are standard error of the mean. **c.** Representative DAB-stained and two immunofluorescent images of a late-stage PD patient using the pS129 α-synuclein antibody (AB_2819037) for images 1 & 2 and a total alpha-synuclein antibody LB509 (AB_2832854) for image 3 showing comparable Lewy pathology produced by the pS129 antibody and increased background in immunofluorescence produced by the total alpha-synuclein antibody. **d.** Detailed aggregate compilation from a late-stage PD patient in immunofluorescence using the pS129 alpha-synuclein antibody (AB_2819037) showcasing characteristic Lewy pathology. **e.** Diffraction-limited aggregate coincidence vs. chance coincidence between pS129 α-synuclein (AB_2270761) and disease-related proteins Ubiqutin (AB_2315524), p62 (AB_398152), TDP-43 (AB_425904), AT8 (AB_223647) and 4R Tau (AB_310014) in a late-stage PD patient. N = 1 **f.** Representative images of Mouse brain tissue imaged on a spinning-disc confocal microscope of SNCA knockout and human SNCA overexpressing mice stained with pS129 antibody (AB_2819037), respectively. **g.** Averaged stack (132 μm x 132 μm x 12 μm) whole-image intensity of mouse brain tissue stained with pS129 antibody (AB_2819037) in SNCA KO mice not expressing α-synuclein and SNCA OVX mice overexpressing human α-synuclein across n = 3 biological replicates per genotype.

**Supplementary Fig. S3.**
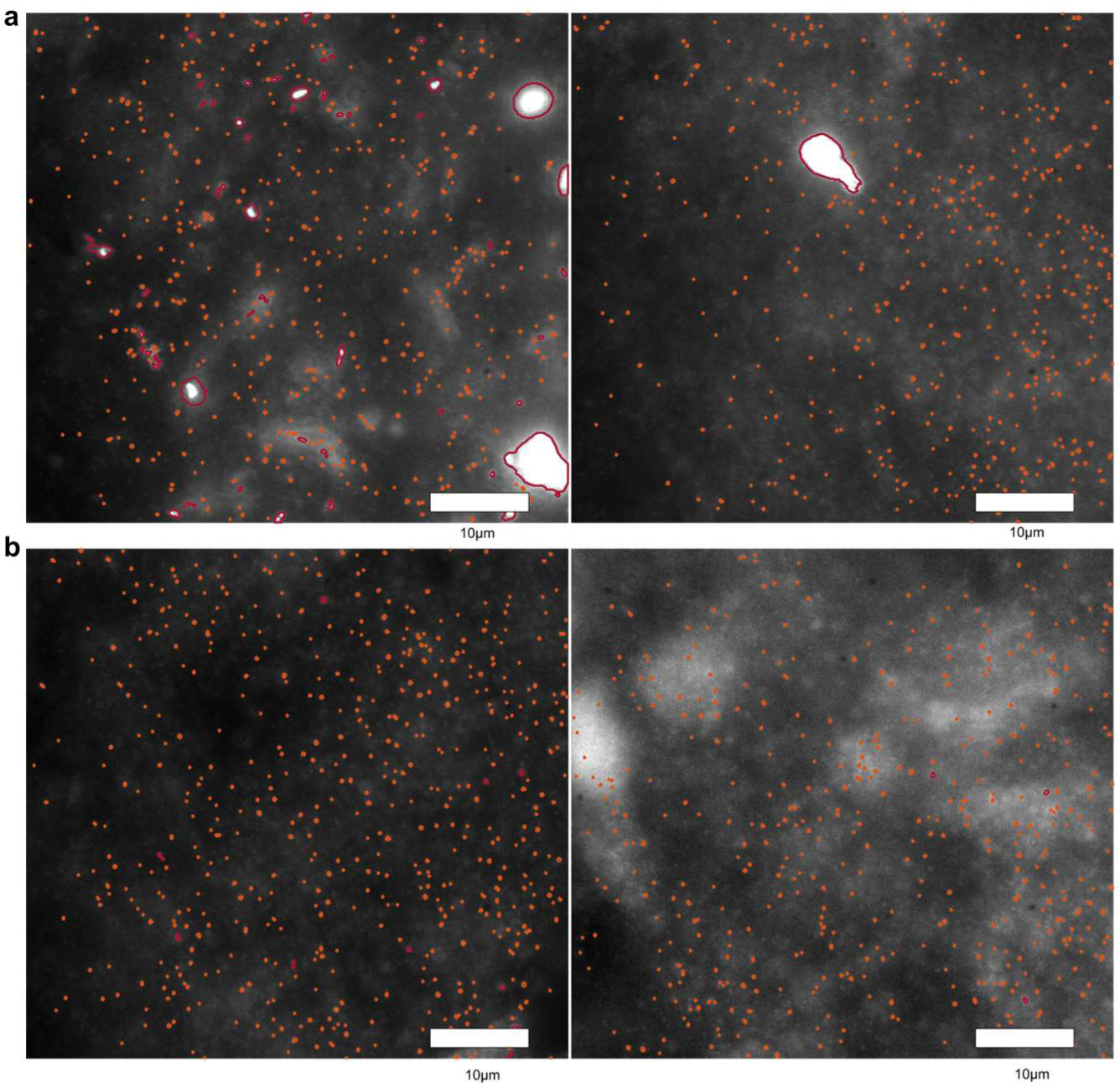
α-synuclein detection and quantification in HC and PD samples. **a.** Representative images of two PD patients stained for pS129 α-synuclein (AB_2270761) showing abundant diffraction-limited puncta. **b.** Representative images of two HCs stained for pS129 α-synuclein (AB_2270761) showing a comparable number of diffraction-limited aggregates detected.

**Supplementary Fig. S4.**
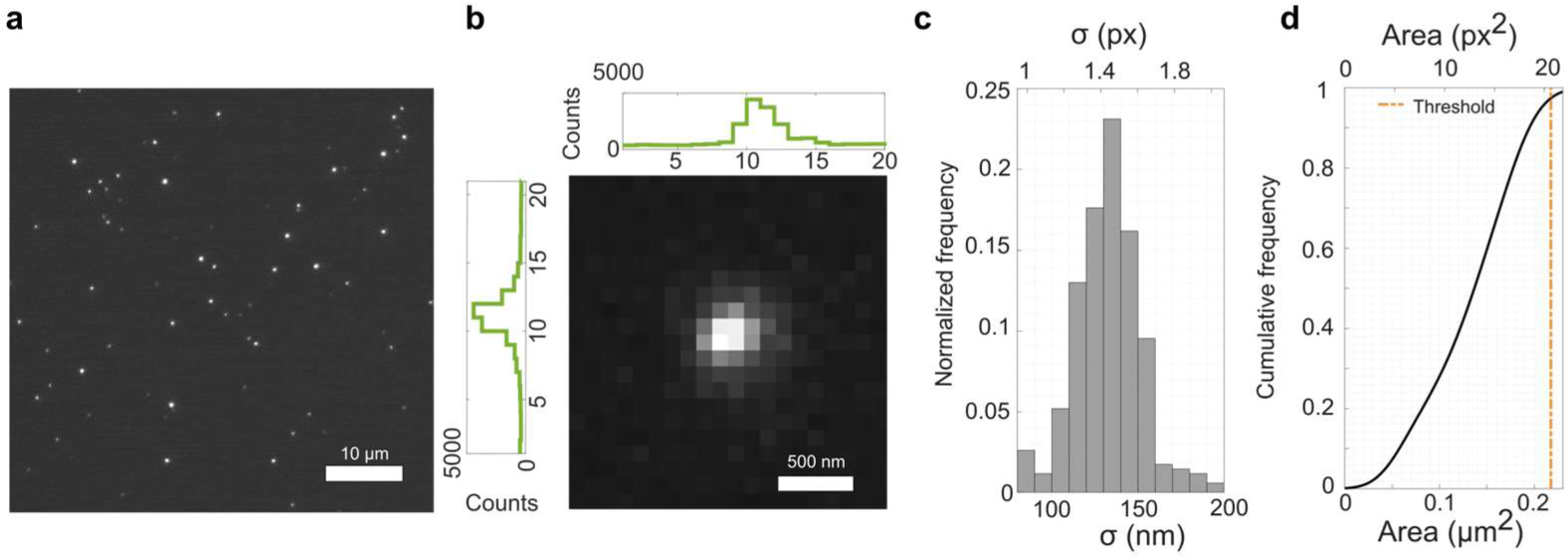
Calibration of the point-spread function with 20 nm fluorescent beads. **a.** Sparse TetraSpeck beads (T7279) were attached to a coverslip with poly-L-lysine and imaged with PBS using 50 mW laser power. **b.** Zoom-in of a single bead with its horizontal and vertical intensity profile. **c.** The corresponding histogram of the width parameter, σ, is extracted by localising each object with a 2D symmetric Gaussian function. **d.** Cumulative frequency of spot sizes was derived using the aggregate detection code with the same parameter used for the human brain data. The cutoff threshold for small aggregates is highlighted in orange.

**Supplementary Fig. S5.**
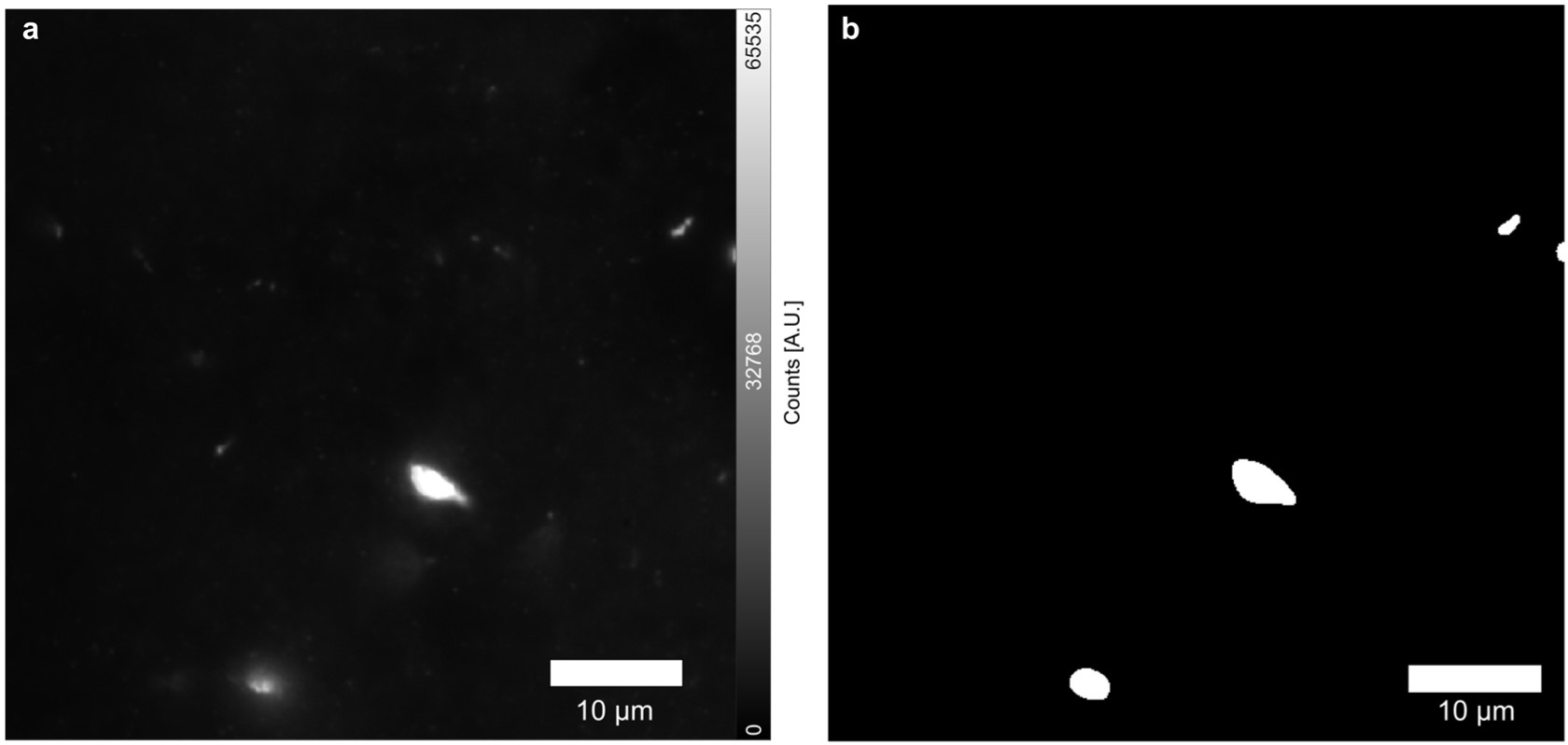
Forming a large-object binary mask. **a.** Original image of a PD patient in the Anterior Cingulate Cortex stained for pS129 α-synuclein (AB_2270761). **b.** Result binary mask from the large object detection code.

**Supplementary Fig. S6.**
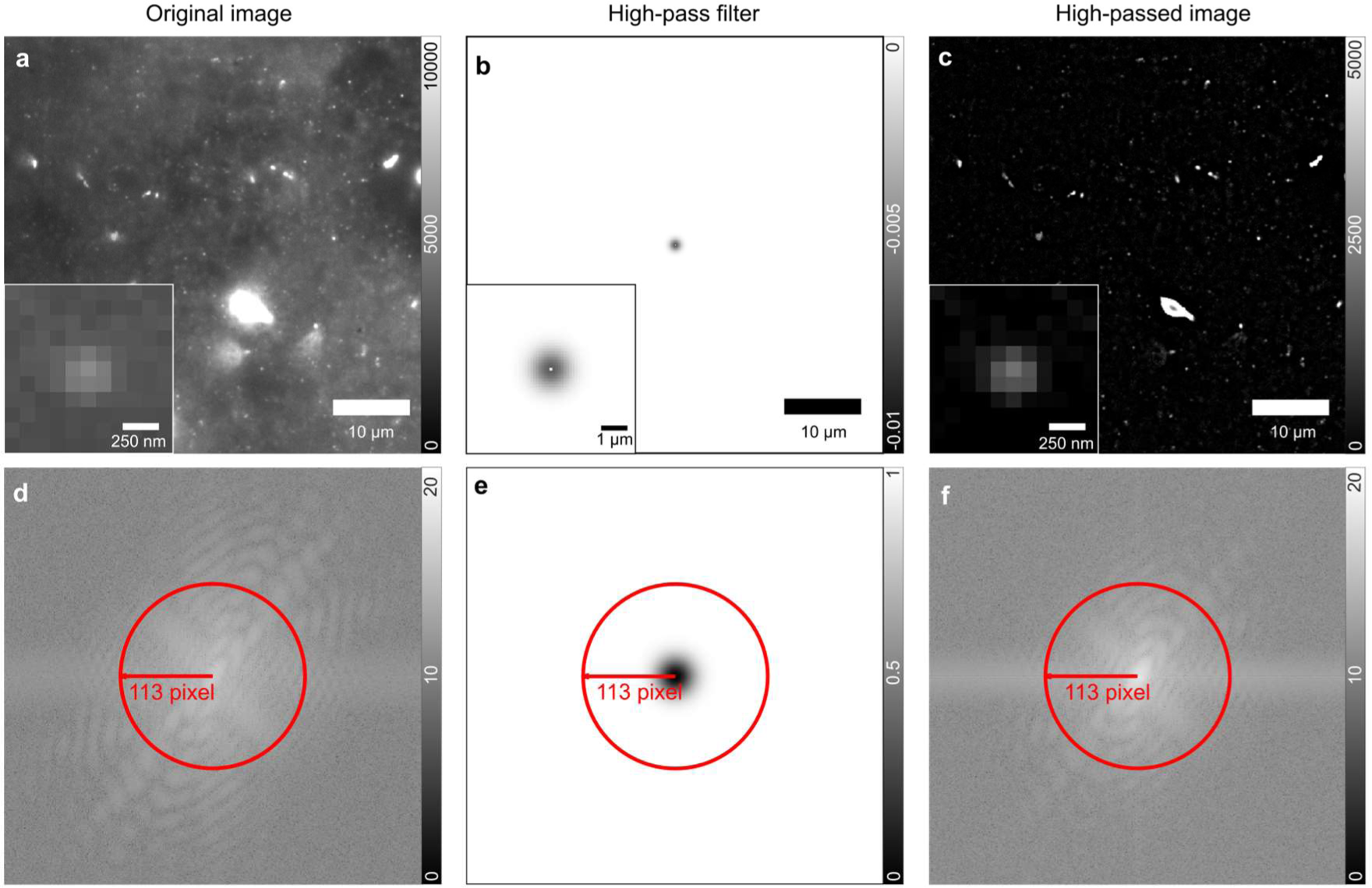
Background removal for oligomer detection. **a-c.** The magnitude of frequency domain of original image, high-pass filter and high-passed image respectively with a highlighted red circle showing the passband due to diffraction limit. **d-f.** The spatial (*i.e.* image. domain of the original image, high-pass filter and high-passed image, respectively. (**a&c)** are shown in log scale.

**Supplementary Fig. S7.**
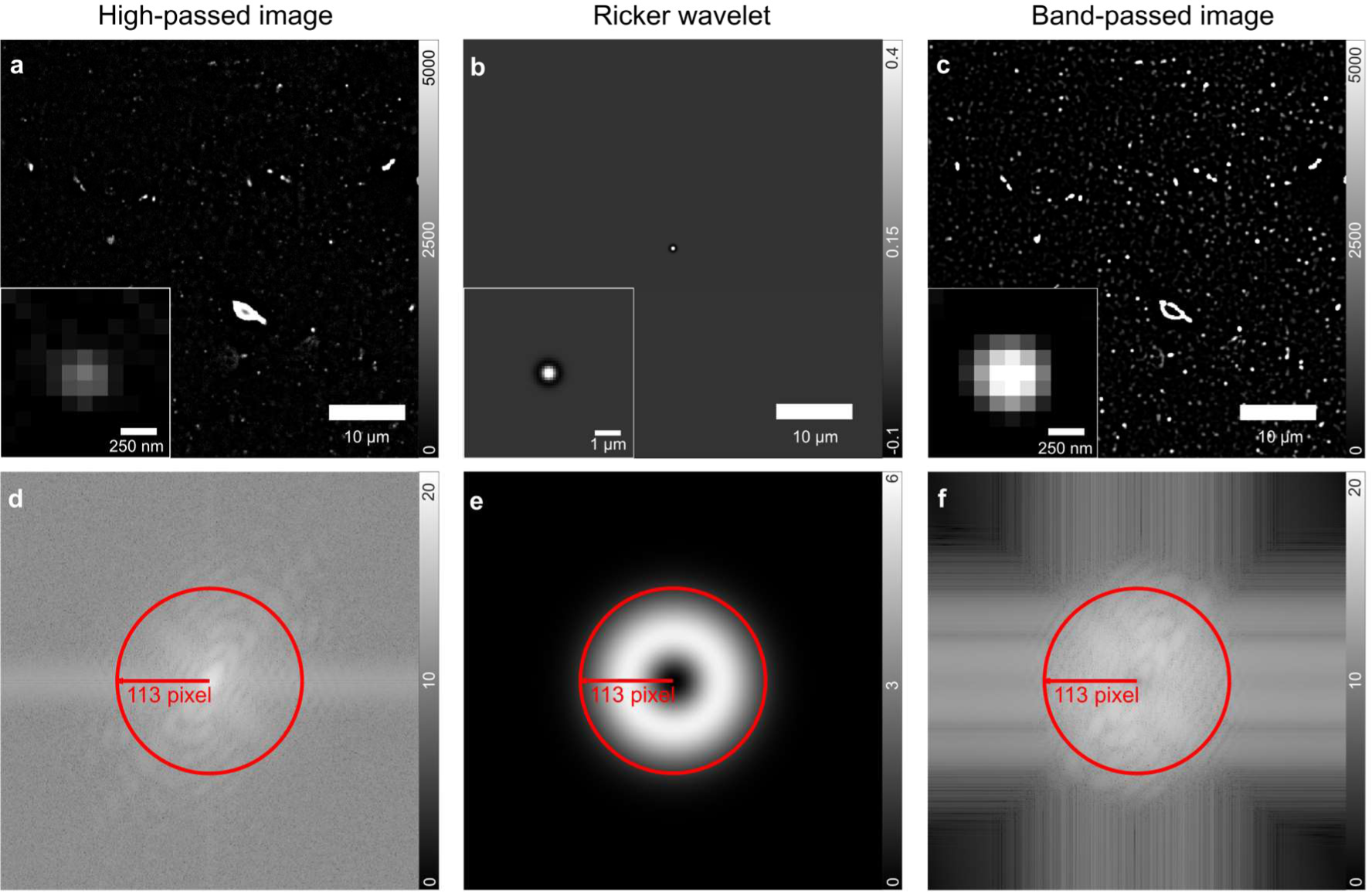
Feature enhancement for oligomer detection. **a-c.** The magnitude of frequency domain of high-passed image, ricker wavelet and band-passed image respectively with a highlighted red circle showing the passband due to diffraction limit. **d.**, **e.**, **f.** The spatial domain of high-passed image, ricker wavelet and band-passed image, respectively. **a.** and **c.** are shown in log scale.

**Supplementary Fig. S8.**
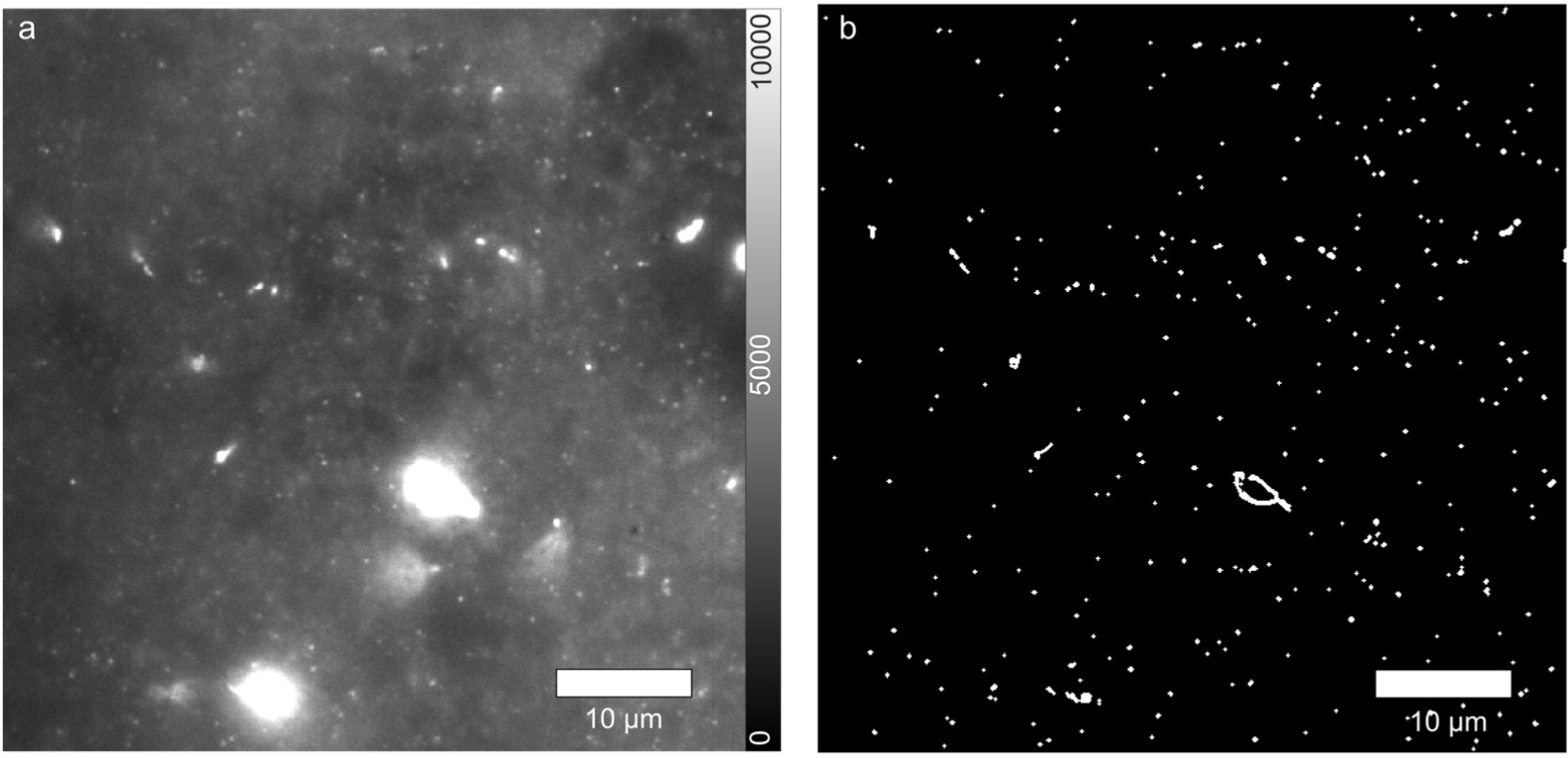
Forming a small-object binary mask. **a.** Original image from the anterior cingulate cortex of a PD sample stained with anti-alpha-synuclein pS129 (AB_2819037) visualised with (AB_144696). **b.** Resulting binary mask after a top 2.5 percentile thresholding.

**Supplementary Fig. S9.**
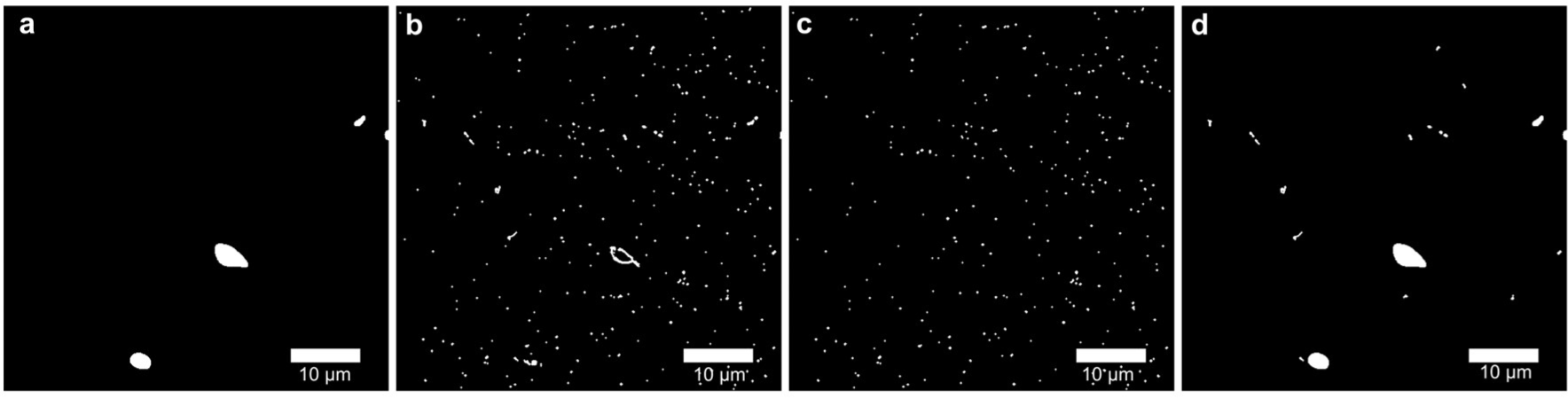
Oligomer and non-oligomer objects after object filtering and classification. **a.** Binary mask from large object detection section. **b.** Binary mask from small object detection session. **c.** Result binary mask for non-oligomer objects **d.** Resulting binary mask for objects classified as diffraction-limited oligomers.

**Supplementary Fig. S10.**
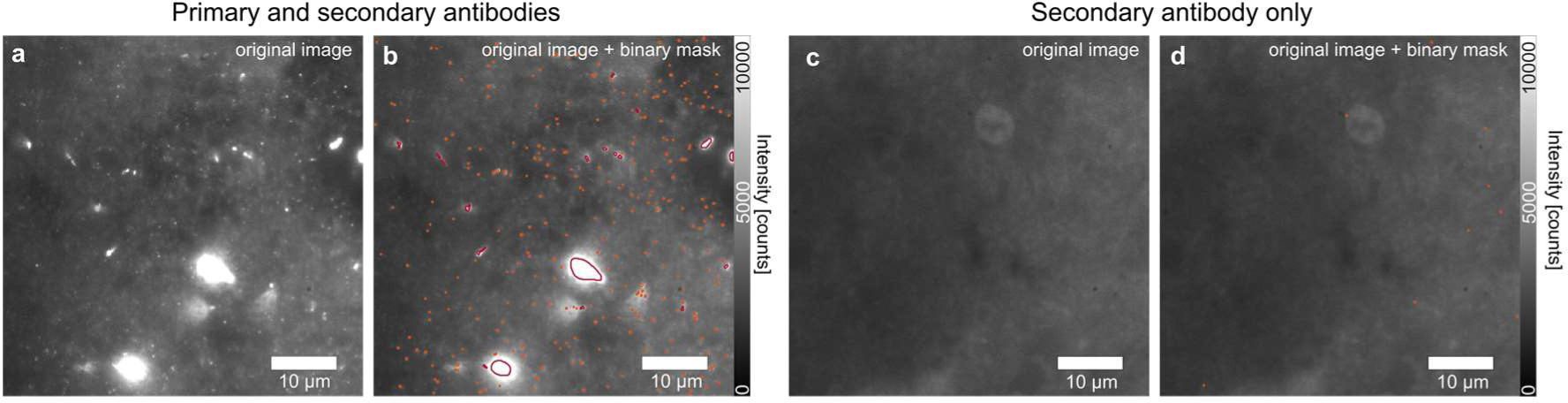
Aggregate-detection output. **a**.**b**. The original image and resulting diffraction limit and non-diffraction limit mask from a PD patient with the primary antibody targeting pS129 alpha-synuclein (AB_2270761) and the secondary Goat anti-mouse AF568 (AB_144696). **c.d.** The original image and resulting diffraction limit and non-diffraction limit mask from a PD patient stained with no primary but only the secondary antibody.

**Supplementary Fig. S11.**
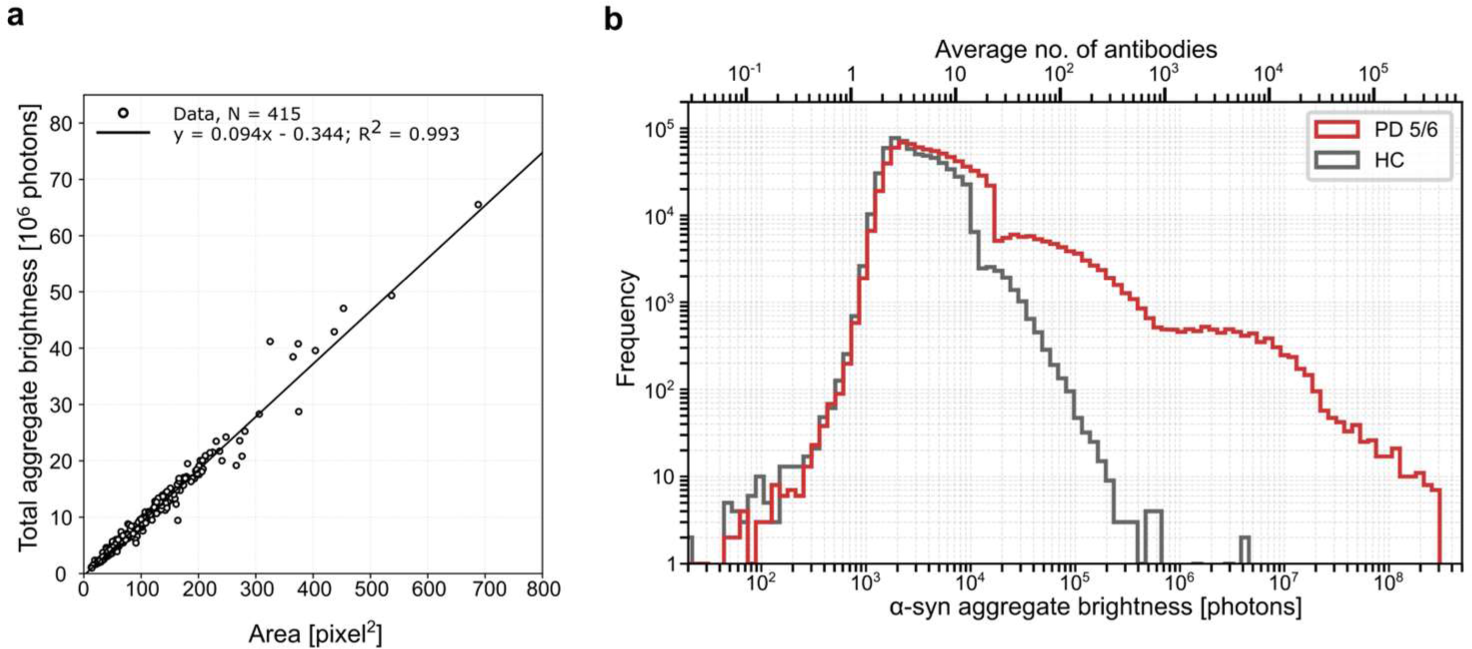
Aggregate brightnesses and size. **a.** Distributions of aggregate brightnesses (photons) plotted against aggregate size (pixels) for *N* = 415 aggregates above the diffraction limit. A linear fit with *R*^2^ = 0.993. **b.** Frequency distribution of the average number of antibodies and the pS129 α-synuclein (AB_2819037) object brightness (photons) for *n* = 910,873 objects detected in PD patients, *n* = 682,364 objects detected in HCs.

**Supplementary Fig. S12:**
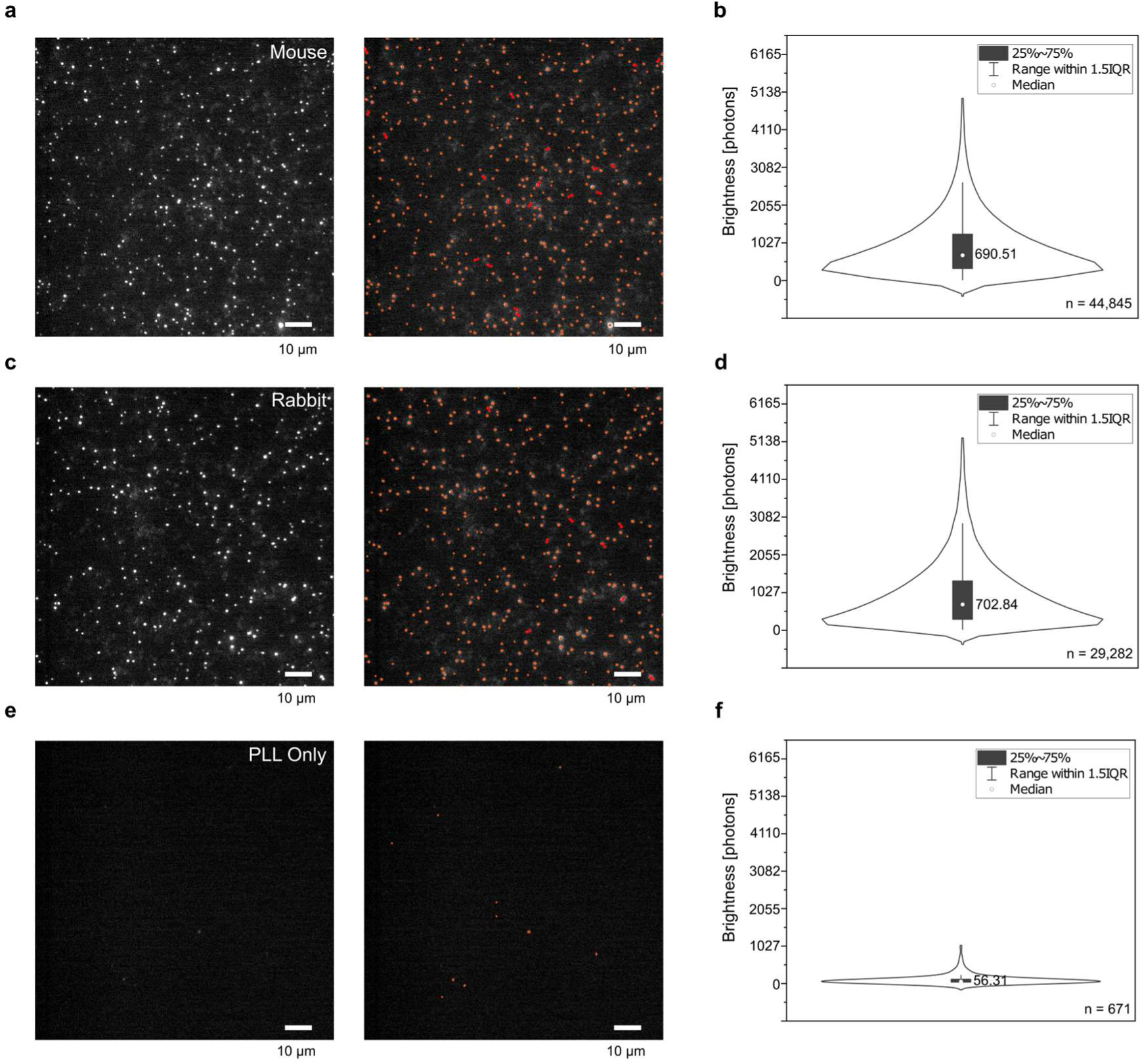
Single-molecule analysis of secondary antibodies. Secondary antibodies (AB_143157, AB_144696) were fixed on a glass coverslip and 50 FoVs across two slides were imaged for each antibody. All imaging conditions were the same as for the main brain imaging. **a.** Representative images of Alexa Fluor 568 goat anti-mouse (AB_144196) and the detection of spots, respectively. **b.** Violin plot of the median brightness = 8,390 of *n* = 44,845 detected spots. **c.** Representative images of Alexa Fluor 568 goat anti-rabbit (AB_143157) and the detection of spots, respectively. **d.** Violin plot of the median brightness = 8,555 of *n* = 29,282 detected spots. **e.** Representative images of negative control (Poly-L-lysine. and the detection of spots, respectively. **f.** Violin plot of the median brightness = 690.5 of *n* = 671 detected spots. Due to the similar incidental signal brightness of these two labelling conditions, we used the mean median brightness of 697 photons as the brightness per antibody under these imaging conditions.

**Supplementary Fig. S13:**
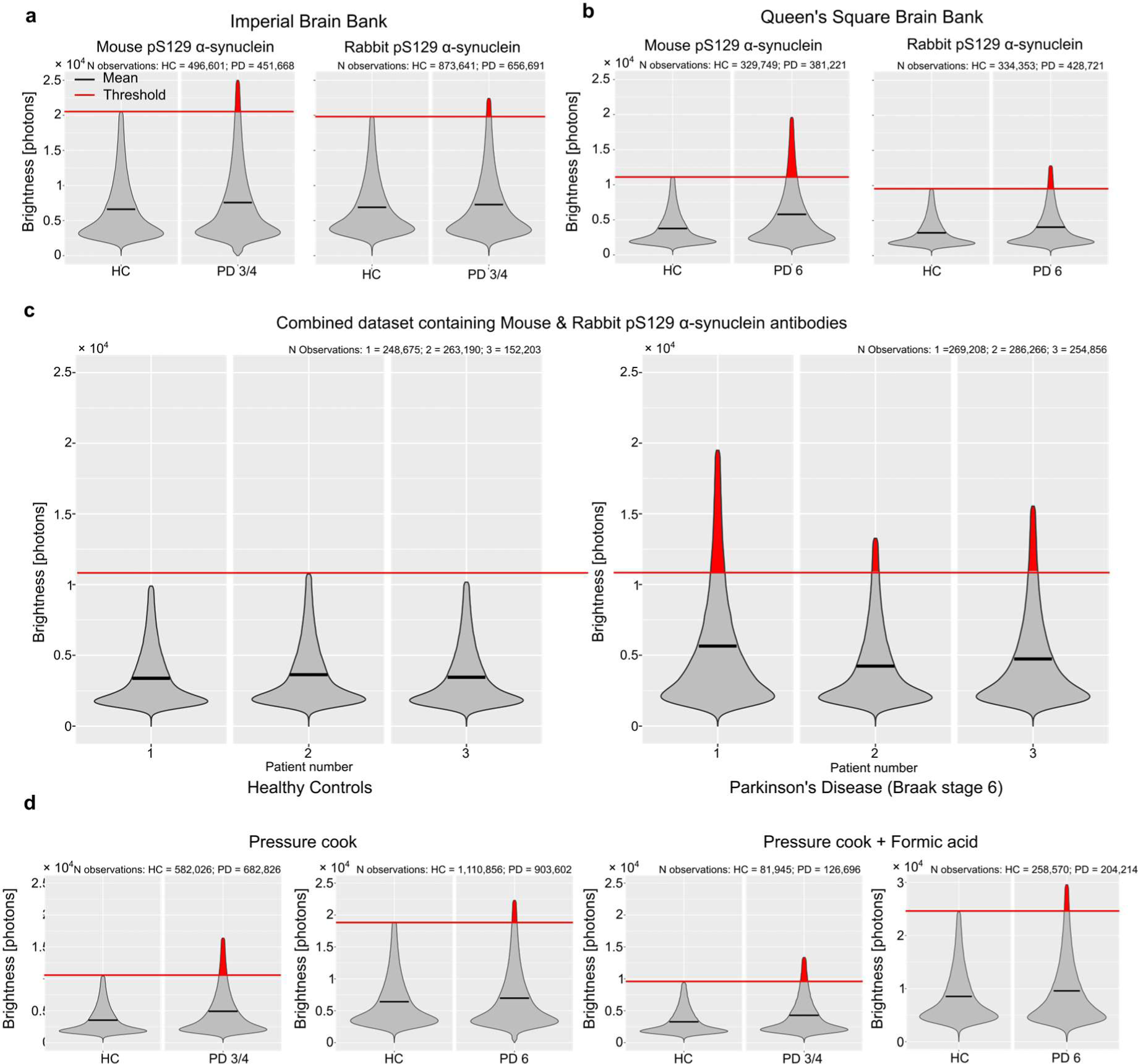
Oligomer brightnesses in PD and HC samples are different across a range of conditions. **a.b.** Violin plots of the intensity (photons) of aggregates detected in *n* = 3 HC and *n* = 3 PD (**a**) Braak stage ¾, and PD (**b**) Braak stage 6) patients using formalin fixation and paraffin embedding (FFPa.. Disease-specific bright aggregates can be detected using the mouse (AB_2819037) and rabbit pS129 alpha-synuclein (AB_2270761) antibodies, respectively, across both mid- and -late-stage disease. **c.** Violin plots of the intensity (photons) of aggregates detected in *n* = 3 HC and *n* = 3 PD (Braak stage 6) patients detected using both alpha-synuclein antibodies (AB_2819037 & AB_2207061). Disease-specific bright aggregates are consistent across all individuals. **d.** Violin plots of the intensity (photons) of aggregates detected in *n* = 3 HC, *n* = 3 PD (Braak stage 3/4) and *n* = 3 PD (Braak stage 6) patients using both alpha-synuclein antibodies (AB_2819037 & AB_2207061). Samples were either treated with a pressure cooker only or pre-treated with formic acid and pressure cooked. Disease-specific aggregates are observed across pre-treatment conditions. All data plots are truncated at 1.5 x IQR.

### 2. Supplementary Tables

**Supplementary Table 1.** Summary of case demographics for human post-mortem brain tissue.

|  | Healthy Control cases | PD, Braak stage 3/4 | PD Braak stage 6 |
| --- | --- | --- | --- |
| Number of cases | 6 | 3 | 3 (+1 neg control only) |
| Brain bank | 3 QSBB, 3 Imperial | Imperial | QSBB |
| Age at death | 87 (71-101) | 80 (73-85) | 75 (73-84) |
| Sex | M | M | M |
| Post-mortem interval [hours]<sup>1</sup> | 45 (18-98) | 21 (16-24) | 68 (38-89) |

<sup>1</sup> Age at death and PMI are given as a mean with range in brackets. PMI - Post-mortem Interval. Negative control is a case that was only used for technical control rather than the main study.

**Supplementary Table 2.** Case Demographics.

| Case | Brain Bank<sup>1</sup> | Pathological diagnosis<sup>1</sup> | Age | Sex | Onset | Duration | PMI (hours) | Confounding pathology | Braak score (α-syn) | Braak and Braak score (tau) | Thal score (amyloid $\beta$) |
| --- | --- | --- | --- | --- | --- | --- | --- | --- | --- | --- | --- |
| 1 | QSBB | PD | 81 | M | 50 | 31 | 73 | mild small vessel disease, microscopic cerebellar infarction, leptomeningeal vessel atheroma | 6 | 2 | 2 |
| 2 | QSBB | PD | 84 | M | 61 | 23 | 71 | mild CAA, hyaline arteriosclerosis, ARTAG | 6 | 1 | 1 |
| 3 | QSBB | PD | 63 | M | 42 | 21 | 38 | mild small vessel disease | 6 | 1 | 0 |
| 4 | QSBB | PD | 73 | M | 62 | 11 | 89 | ARTAG, microinfarcts, CAA, hyaline arteriosclerosis, atherosclerosis | 6 | 1 | 3 |
| 5 | QSBB | Control | 87 | M | - | - | 40 | moderate small vessel disease | 0 | 1 | 0 |
| 6 | QSBB | Control | 101 | M | - | - | 60 | ARTAG, CAA, hyaline arteriosclerosis, atherosclerosis | 0 | 1 | 0 |
| 7 | QSBB | Control | 88 | M | - | - | 98 | Pathological ageing, CAA | 0 | 2 | 2 |
| 8 | Imperial | PD | 73 | M |  |  | 24 |  | 3 | 1 | 3 |
| 9 | Imperial | PD | 81 | M |  |  | 22 |  | 4 | 2 |  |
| 10 | Imperial | PD | 85 | M | 70 | 15 | 16 |  | 4 | 2 |  |
| 11 | Imperial | Control | 71 | M | - | - | 29 | Pathological ageing | 0 | - | - |
| 12 | Imperial | Control | 91 | M | - | - | 18 | PART, ARTAG | 0 | - | - |
| 13 | Imperial | Control | 85 | M | - | - | 23 | Pathological ageing | 0 | - | 3 |

<sup>1</sup>PD, Parkinson's disease; QSBB, Queen's Square Brain Bank; PMI, Post-mortem Interval

**Supplementary Table 3.**
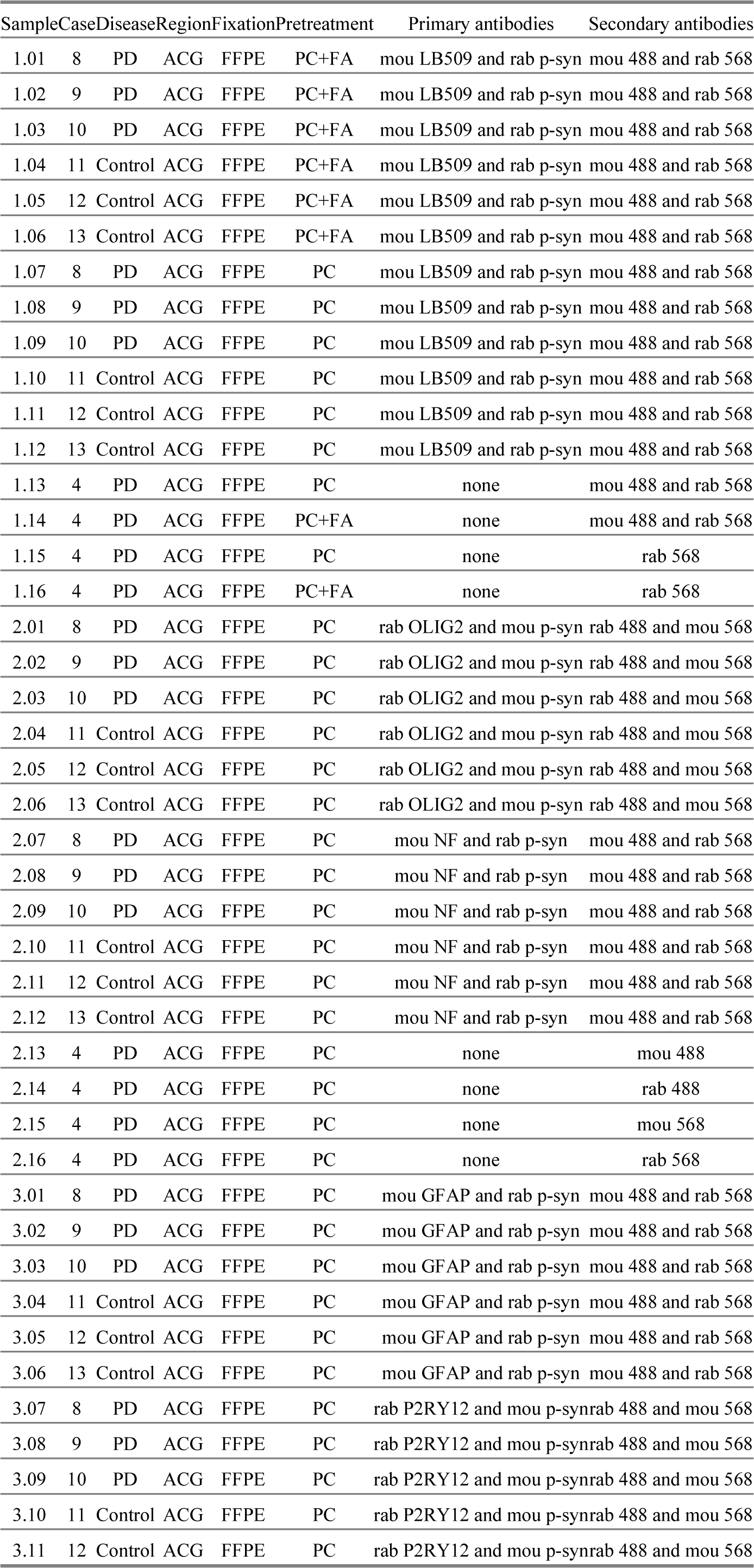
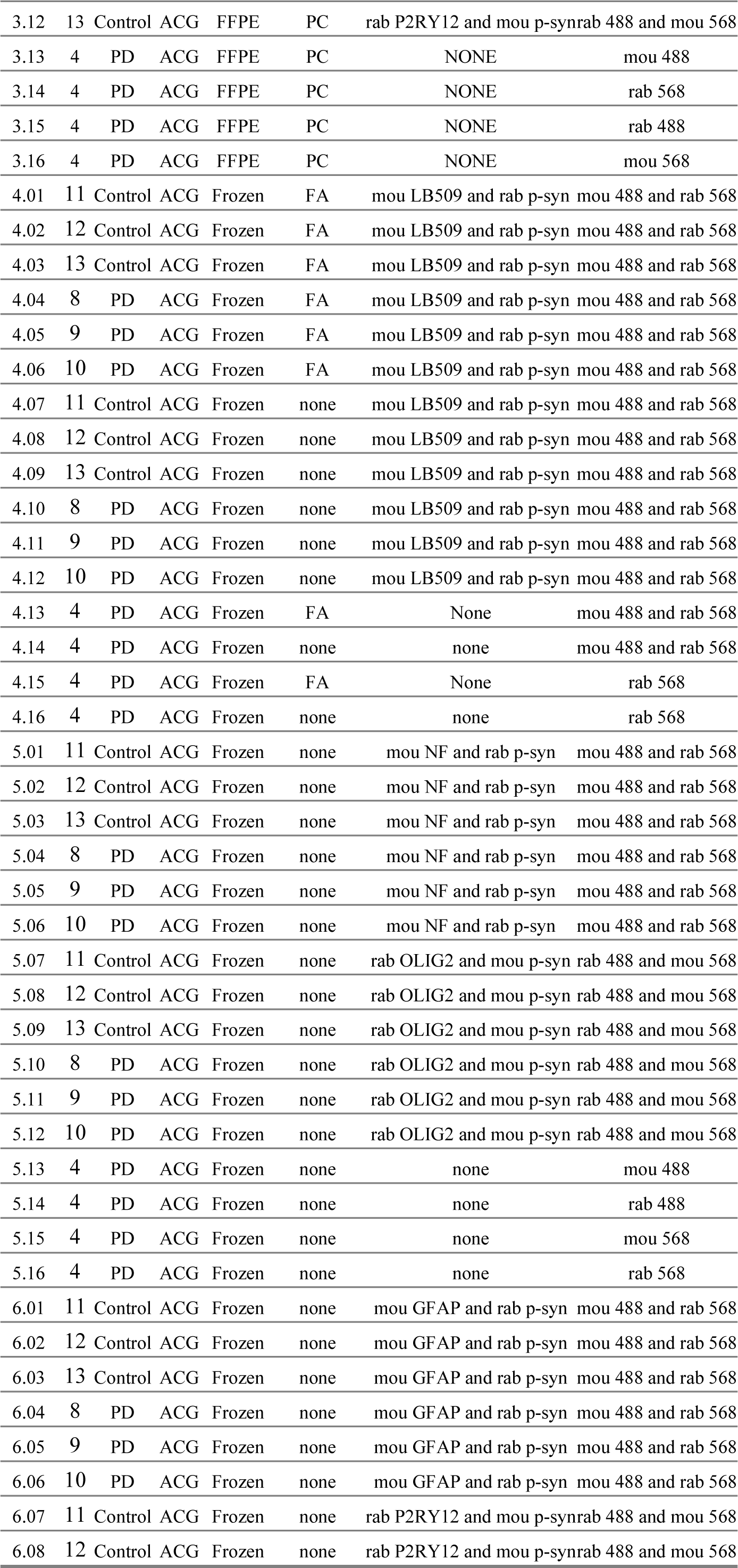
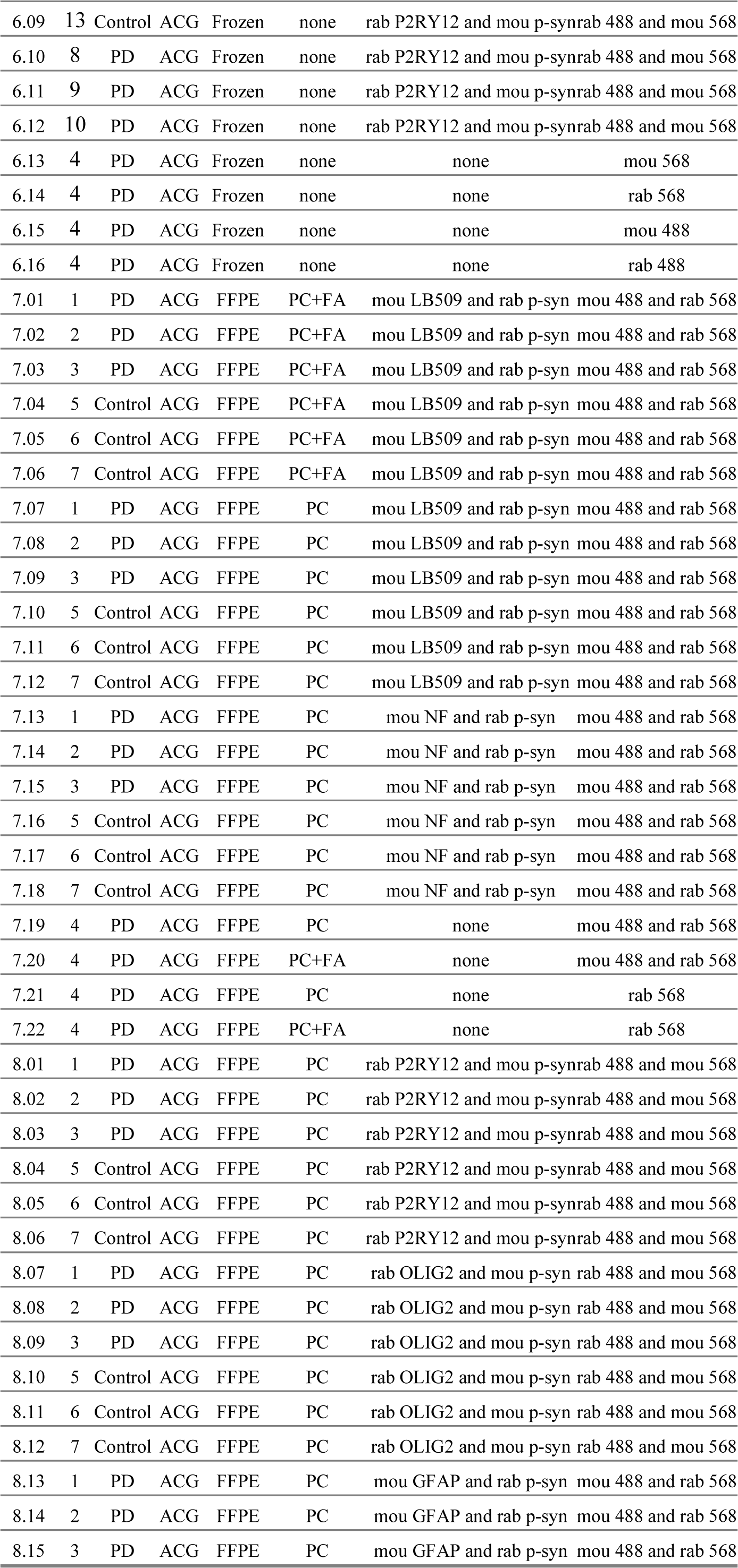
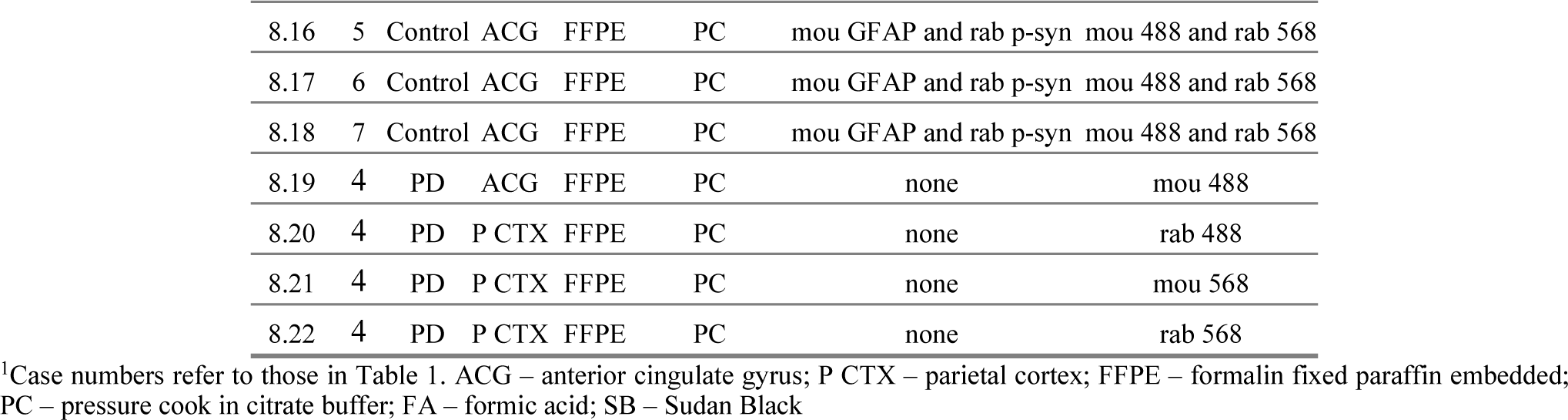
Staining Plan Table.

**Supplementary Table 4.**
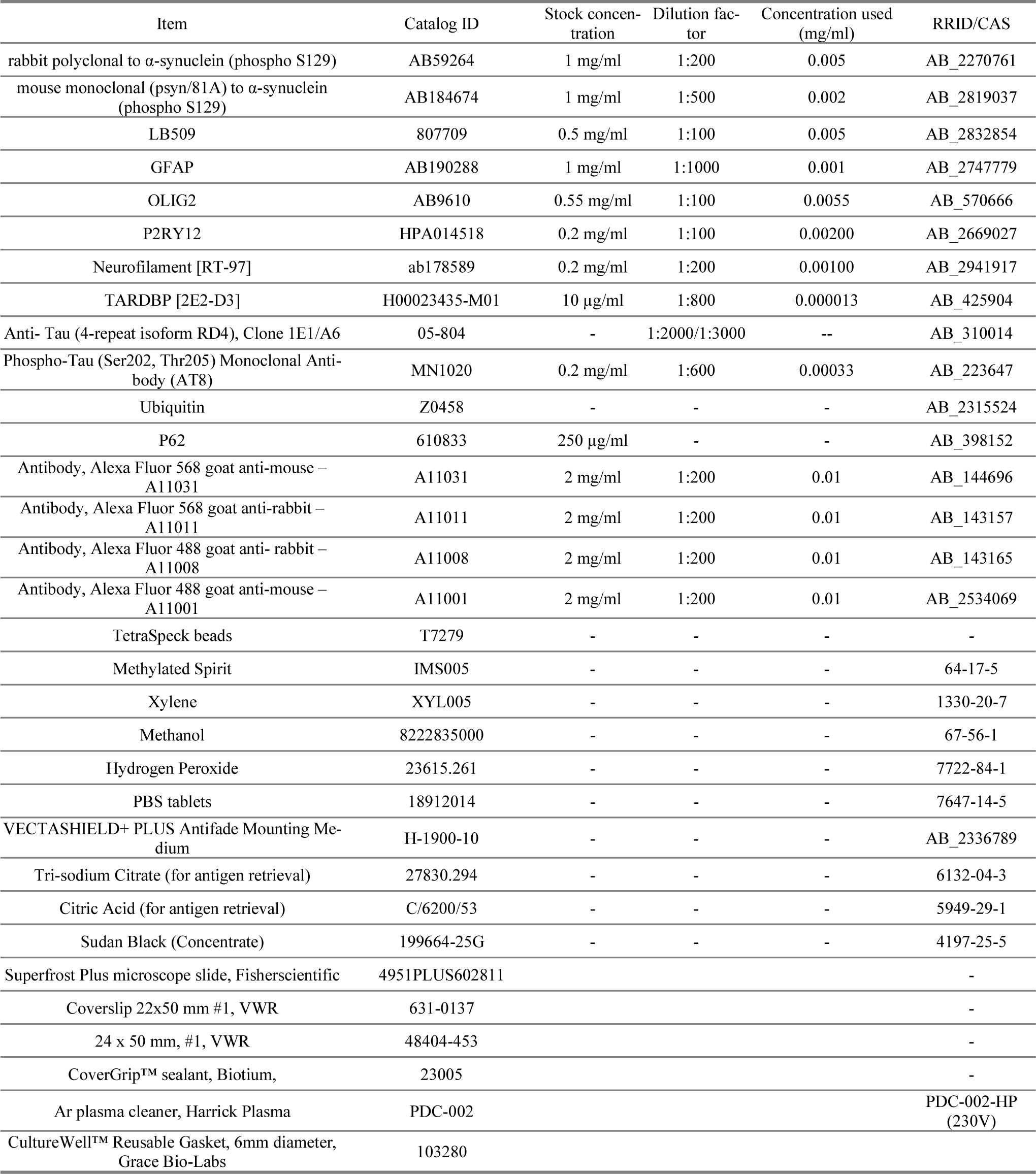
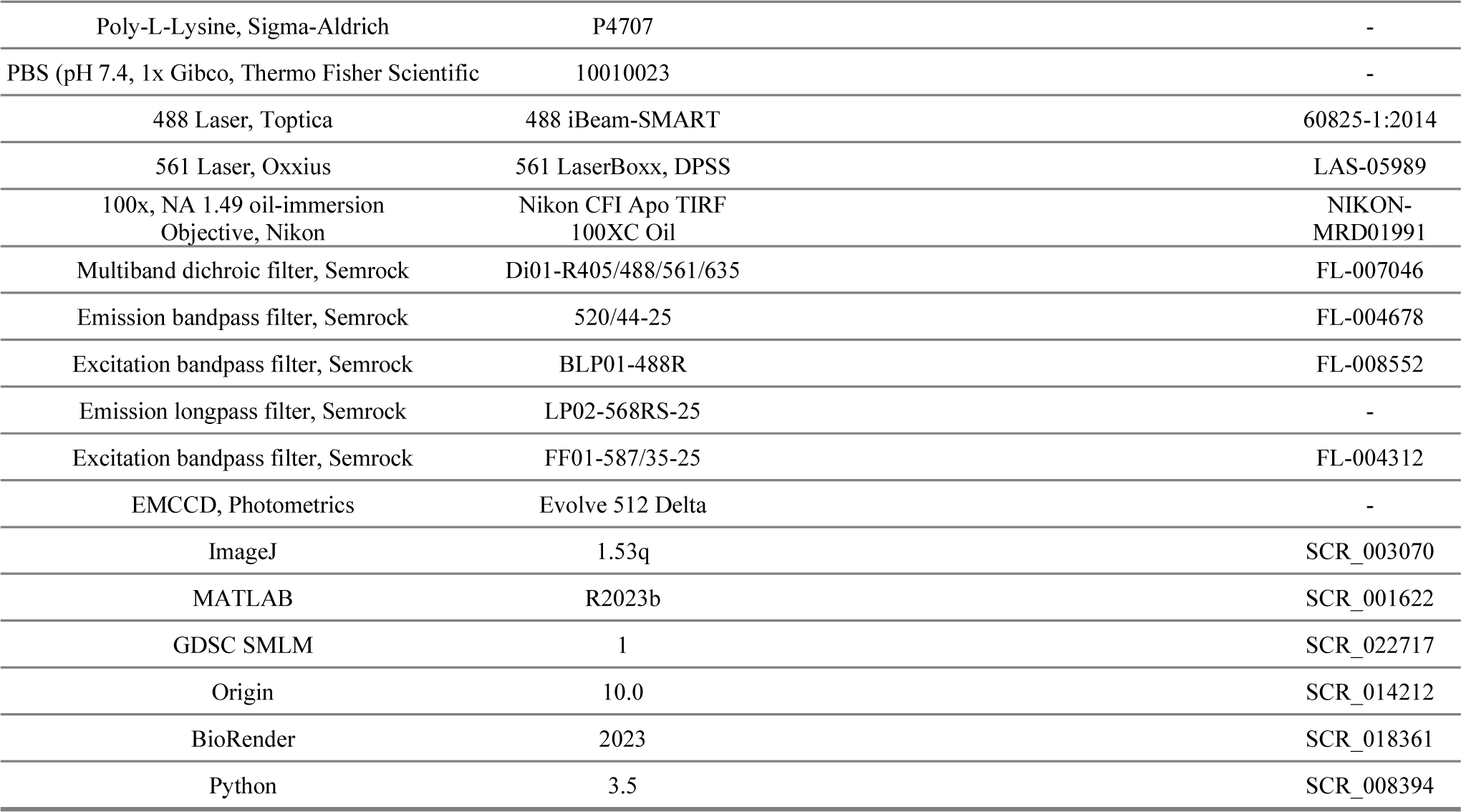
Reagents and tools.

### 3. Supplementary Note 1: Aggregate-detection pipeline

#### Characterisation of system point spread function

It is important to distinguish nanoscale objects (oligomers), whose appearance on a detector is dominated by the diffraction of light, from objects larger than the diffraction limit (*i.e.* medium-sized aggregates, LBs and LNs). Even though fluorescently labelled nanoscale emitters appear approximately identical, slight variations exist due to out-of-focus emitters, optical aberrations, background variance and other sources of noise (*e.g.* shot noise and read noise). To characterise the measured area for sub-diffraction objects, images of beads (40 nm, FluoSphere F10720) were analysed with our pipeline (Supplementary Figure S4). As very few beads are defocused and/or aggregated (*i.e.* larger than the diffraction limit), the area threshold is selected to include 97% of all data. This threshold then was then applied to classify the detected objects into oligomer and non-oligomer categories.

#### Step1 - Large object detection

The hallmark of Parkinson’s disease is the presence of large aggregates of α-synuclein such as Lewy Bodies and Lewy Neurites. These comparatively very bright objects mask the presence of much dimmer nanoscale objects, such as oligomers, and it is therefore necessary to exclude regions of the image containing these and other large features. To do so, a large-object mask is made using a difference-of-Gaussians kernel, *i.e.* a band-pass in the frequency domain to eliminate both the background and small objects (Supplementary Equation S1, where σ_1_ is 2 pixels, slightly larger than σ of the diffraction limit, and σ_2_ uses 40 pixels, larger than most of LBs and LNs thus eliminating the background as much as possible). Where the large objects to be detected are instead cell types, σs are: **microglia**, σ_1_ = 2 pixels, σ_2_ = 10 pixels; **neurons & astrocyte**: σ_1_ = 0.5 pixels, σ_2_ = 5 pixels; **oligodendrocyte**: σ_1_ = 4 pixels, σ_2_ = 40 pixels.

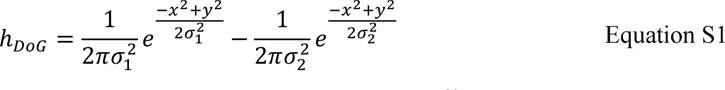

Since the area of LBs is large compared to the pixel size, a two-class Otsu’s threshold,^83^ which separates pixels into two classes, is then applied. The resulting binary classification on an example image is shown in Supplementary Figure S5.

#### Step2 - Image filtering

Two different frequency-filtering steps are applied to remove the background and emphasise the diffraction-limited spots in the image, i.e., the oligomers. In this dataset, the background due to autofluorescence of the brain tissue was relatively smooth. It can therefore be regarded as a low-frequency signal suppressible with a high-pass filter. This was implemented by subtracting the low-pass filtered image from the original image. Supplementary Equation S2 describes the spatial representation of the high-pass kernel, where δ is the Dirac pulse at (x,y) = (0,0), σ = 5 pixel larger than σ from sub-diffraction beads (1.3 pixel), and designed to maintain small and intermediate size aggregates that were the focus of our study. The original image, high-pass kernel and high-passed image are shown in Supplementary Figure S6.

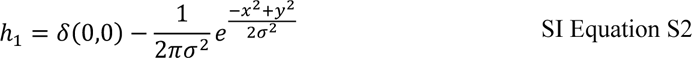

After removing the low-frequency autofluorescence background, high-frequency pixel noise hinders object detection. To improve the detectability of these small objects, a feature enhancement filter, namely the Ricker wavelet is used.^46^ This step acts as a band-pass filter. The wavelet, described in Supplementary Equation S3 uses σ = 1 pixel such that it is slightly smaller than the σ = 1.3 pixel from sub-diffraction beads to maximise the suppression of pixel noise. Supplementary Figure S7 shows the spatial and frequency representation of the wavelet and band-pass image, respectively. The overall formula is described in Supplementary Equation S4, where H_1_ and H_2_ represents the Fourier transform of h_1_ and h_2_ and i_init_ and i_filtered_ represents the image in the spatial domain. The system bandwidth, calculated in pixels in Supplemental Equation S5 as the Rayleigh diffraction limit in the frequency domain, is shown in Supplementary Figures S8 and S9 as the red circle.

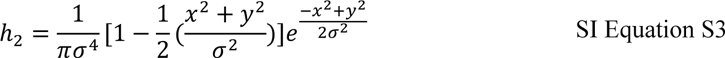

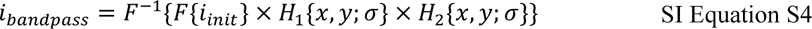

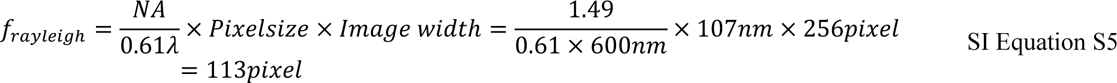

#### Step3 - Image thresholding

After background and autofluorescence suppression, a threshold is used to identify potential regions of interest within the image. Due to the variability in the number of diffraction-limited spots visible during imaging (typically between ∼30– 300), a threshold value of top 2.5 percentile of the filtered image value was used. This value was determined experimentally based on the relatively small percentage of pixels containing oligomers, while avoiding the influence of a small number of very bright objects that sometimes remained after the previous filtering steps. The resulting binary mask is shown in Supplementary Figure S10.

#### Step 4 - Object filtering and classification

Due to the diffraction limit of light, nanoscale fluorescent objects are diffraction-limited and appear to have similar shapes. A morphology filter is used to reduce the number of false positives due to spuriously bright pixels. Specifically, an opening filter is applied by first eroding and then dilating the binary image with a disk shape with a diameter of 3 pixels. Furthermore, there are also false positives from LBs in the FoV, which can be eliminated by the known position from the large object detection section. Supplementary Equation S6 represents the object filtering process where *M_binary* is the binary mask from the large object detection and B_struct is a pre-defined binary structure, specifically a binary disk with radius 1. This effectively compares every object in the image with the disk structure, and everything smaller than the disk or dissimilar to the disk will be erased. Further details can be found in Ref^84^. Objects are then classified into oligomer and non-oligomer objects separately.

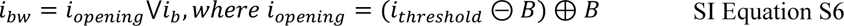

### 4. Supplementary Note 2: Simulations

In the following section, we describe the modelling assumptions and calculations done to simulate the expected effective signal-to-noise ratio of imaging single bright objects in a noisy tissue background.

We model the expected numbers of photons from a signal object and from a fixed concentration of background objects, emitting in the illuminated volume of one pixel. The pixel axial length was assumed as a depth of field is

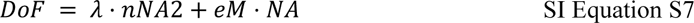

where λ is the wavelength of excitation light (561 nm here), *n* the refractive index of the medium (assumed 1.515), NA is the numerical aperture, *e* is the smallest resolvable object on the camera plane (11 µm here) and M is the magnification (assumed 100 here).

We assume one bright single object that emits in this pixel, and background objects at a concentration of 100 µM (the number in the pixel will vary, dependent on the NA-dependent depth of field). We assume each background object has a photon emission rate of 122 photons s-1—this number was chosen to agree with experiment in this work. We assume that the signal objects may have one of two brightness levels, dependent on if they are “large objects” or “small objects” observed in experiment. For small objects we assume a photon emission rate of 13,000–21,000 photons s-1 and for large objects we assume a photon emission rate of between 24,000–300,000 photons s-1. Again, we emphasise that these values were chosen to agree with experiment.

Thus, the number of photons arriving from our single signal object is set, and the number of background photons is calculated in a numerical-aperture dependent manner by multiplying the volume (DoF*107^2^ nm^2^) by the concentration of background objects.

We then assume that the numerical aperture and the camera quantum efficiency will cause a loss of photon detection efficiency, and therefore a loss in the number of detected photons. The numerical aperture will cause a loss of efficiency proportional to the cone angle of collected fluorescence:

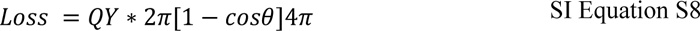

where the angle, θ, is calculated from the numerical aperture by Supplemental Equation S9.

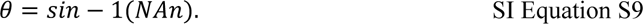

e then, assuming a fixed numerical aperture, calculate the signal-to-noise ratio (effectively as in Moerner and Fromm^85^) by calculating the number of signal photons detected in 100 ms, number of background photons detected in 100 ms, and calculating the signal-to-noise ratio as:

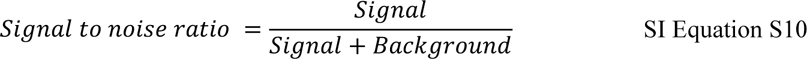

## Notes

https://doi.org/10.5281/zenodo.10610657

https://doi.org/10.5281/zenodo.10610924

https://doi.org/10.5281/ZENODO.8319436

